# The chromosome-scale genome of the drought-adapted Caatinga tree *Spondias tuberosa* reveals a conserved, unduplicated architecture within Anacardiaceae

**DOI:** 10.64898/2026.11.10.765701

**Authors:** Cícero Almeida, Ellen Lopes, Daniel Santos, Gustavo Rodrigues, José Vieira Silva, Aaryan Bhatia, André Marques

## Abstract

**Background and Aims:** *Spondias tuberosa* (umbu) is a tree endemic to the semi-arid Caatinga of north-eastern Brazil, adapted to recurrent drought and high temperatures. Although genomes of mango (*Mangifera indica*), pistachio (*Pistacia vera*) and cashew (*Anacardium occidentale*) are available, comparative analyses of genome evolution across Anacardiaceae remain scarce. We aimed to generate a chromosome-scale reference genome for *S. tuberosa* and use it to examine genome architecture, meiotic recombination and gene family evolution within the family.

**Methods:** We assembled the genome of a single individual using PacBio HiFi and Omni-C (Hi-C) data, mapped meiotic crossovers by single-nucleus RNA sequencing of pollen, and compared the genome with those of six other Sapindales species through orthology, synteny, synonymous substitution (Ks) and gene family expansion/contraction analyses within a dated phylogenomic framework.

**Key Results:** The assembly anchors 288 Mb onto 16 chromosomes (98.8% BUSCO completeness) and contains 24,232 protein-coding genes, 52.01% repetitive DNA and 1.78 million heterozygous SNPs (0.62%). We identified 1,740 crossovers in 153 pollen nuclei (4.01 cM Mb^−1^), which were depleted in repeat-rich pericentromeric regions. Ks analyses revealed a whole-genome duplication shared by *Mangifera* and *Anacardium* that is absent from *S. tuberosa*, which retains only the ancient γ triplication. The 97 gene families expanded in *S. tuberosa*, mostly through tandem duplication, include receptor-like kinases, glutathione *S*-transferases and disease-resistance proteins, consistent with roles in environmental sensing and stress tolerance.

**Conclusions:** Genome structure has remained highly conserved across Sapindales, except in the mango–cashew lineage, which underwent extensive rearrangements following a whole-genome duplication. As sister to the remaining Anacardiaceae sampled and with an unduplicated, largely collinear architecture, *S. tuberosa* provides a reference for comparative mapping and for transferring genomic information to economically important woody relatives.

## INTRODUCTION

Tree breeding programs face challenges that distinguish them from annual crop improvement, largely due to the protracted juvenile phase and long reproductive cycles of woody perennials, which delay the evaluation of mature traits and often require decades per breeding cycle (Grattapaglia *et al*., 2018). These constraints are compounded by high heterozygosity, pronounced inbreeding depression, and the scarcity of high-quality reference genomes for non-model species (Harfouche *et al*., 2019; Hickey *et al*., 2019), limiting the application of modern strategies such as genomic selection under increasingly unpredictable genotype-by-environment interactions (Cooper *et al*., 2021).

The genus *Spondias* (Anacardiaceae) comprises c. 18 species distributed in tropical America and Asia, several of which are of economic importance owing to their fruits (Mitchell and Daly, 2015), including *S. mombin* L., *S. dulcis* Parkinson, *S. purpurea* L., *S. bahiensis*, and *S. tuberosa* Arruda Câmara, all sharing a similar karyotype (2n = 32; Almeida *et al*., 2007). *S. tuberosa* occurs in the Caatinga, a seasonally dry tropical forest biome in northeastern Brazil marked by low precipitation (300–800 mm yr⁻¹) and high temperatures (27–29 °C) (Balbino *et al*., 2018). The species is andromonoecious, bee-pollinated, and self-incompatible (Nadia *et al*., 2007), and its fruits are widely consumed for their nutritional and antioxidant properties (Dias *et al*., 2019; Rodrigues *et al*., 2024).

High-quality genome assemblies are essential for advancing functional genomics, evolutionary biology, and stress-adaptation research (Bernal-Gallardo and de Folter, 2024; Varshney *et al*., 2024). Within Anacardiaceae, such assemblies are already available for *Mangifera indica* (Wang *et al*., 2020; Bally *et al*., 2021; Ma *et al*., 2021; Singh *et al*., 2021; Wijesundara *et al*., 2024), *Toxicodendron vernicifluum* (Bai *et al*., 2022), and *Pistacia vera* (Kafkas *et al*., 2023), and a draft genome and transcriptomic resources are available for

*Anacardium occidentale* (Capelari *et al*., 2021; Savadi *et al*., 2022). Genome sizes within the family are moderate, ranging from approximately 290–375 Mb in *M. indica* to 623.3 Mb in *P. vera*, with repetitive content between 34.9% and 61.7% (Wang *et al*., 2020; Bai *et al*., 2022). Despite this growing genomic resource base, comparative studies exploring evolutionary relationships and synteny across the family remain limited.

Because *S. tuberosa* is native to semi-arid environments, its genome offers an opportunity to better understand plant strategies for coping with water deficit, as well as aspects of its reproductive biology. Here, we report a chromosome-scale genome assembly of *S. tuberosa* generated using PacBio HiFi and Hi-C data, enabling gene prediction, characterization of repetitive DNA, construction of a genome-wide crossover landscape from single pollen nuclei, analysis of gene family expansion and contraction, and investigation of synteny and whole-genome duplication history within Anacardiaceae. We hypothesize that the genome of *S. tuberosa* harbors structural and functional features — including gene family expansions — associated with adaptation to the drought-prone conditions of the Caatinga biome, and that, as a member of subfamily Spondioideae — sister to the remaining Anacardiaceae sampled — it retains a highly conserved genomic architecture, positioning it as a valuable reference for comparative genomics across the family.

## MATERIALS AND METHODS

### Plant material, DNA isolation, and sequencing

Leaves were collected from a single adult individual of *Spondias tuberosa* growing in a natural population in northeastern Brazil (Arapiraca, Alagoas State). The same individual was used for all sequencing datasets generated in this study (PacBio HiFi, Omni-C, Illumina paired-end and mate-pair, and pollen single-nucleus RNA-seq), which is required for the haplotype-based detection of crossovers. High-molecular-weight DNA was extracted from 1.5 g of fresh leaf tissue using the NucleoBond HMW DNA kit (Macherey-Nagel). DNA quality was assessed using a FEMTO Pulse system (Agilent), and concentration was measured with a Quantus fluorometer (Promega). A PacBio HiFi library was prepared following the manufacturer’s protocol (“Preparing HiFi SMRTbell® Libraries using SMRTbell Express Template Prep Kit 2.0”). DNA fragmentation was performed using g-Tubes (Covaris), and size selection was carried out using the SageELF system (Sage Science). Library size distribution was verified using FEMTO Pulse (Agilent). Sequencing was performed on the Sequel II platform using the Sequel II Binding Kit 2.0 and Sequencing Kit 2.0 with a 30-hour movie (Pacific Biosciences).

For Omni-C (a sequence-independent Hi-C method) library preparation, chromatin was extracted from 0.5 g of fresh leaf tissue, and library construction followed the manufacturer’s recommendations for plant samples (Dovetail Genomics). The final Illumina-compatible library was sequenced in paired-end mode (2 × 150 bp) on a HiSeq 3000 platform (Illumina).

For short-read sequencing, genomic DNA was fragmented to 400–600 bp, and libraries were prepared using the Nextera DNA Sample Preparation Kit (Illumina), following the manufacturer’s instructions. Sequencing was performed on the Illumina HiSeq 2500 platform (paired-end 2 × 100 bp). A mate-pair library was prepared from high-molecular-weight genomic DNA using the Nextera Mate Pair Sample Preparation Kit (Illumina), following the manufacturer’s instructions for gel-free preparation with an insert size > 2 kb, and was sequenced on the Illumina HiSeq 2500 platform. All sequencing was carried out at the Central Laboratory for High Performance Technologies in Life Sciences (LacTad), University of Campinas (UNICAMP), Brazil, and GenOne Tecnologia (www.genone.com.br).

### Genome assembly

HiFi reads were assembled using hifiasm (Cheng *et al*., 2021) with Hi-C integration, generating a primary assembly along with two fully phased haplotype assemblies (haplotype 1 and haplotype 2). Contigs were then scaffolded into chromosome-scale pseudomolecules with HapHiC (Zeng *et al*., 2024) using the Omni-C reads. The primary assembly was used as the reference genome for all downstream analyses (annotation, synteny, SNP calling, gene family analyses, and genome feature visualization), except for crossover identification, for which haplotype 1 was used as the reference (see ’Crossover identification’ below).

### Genome annotation

Microsatellite sequences were identified using GMATA (Wang and Wang, 2016), considering di-to heptanucleotide motifs with a minimum of five repeat units. Satellite DNA arrays were annotated by mapping previously described satellite monomers of *S. tuberosa* (Martins *et al*., 2025) onto the assembly in Geneious R9, and the resulting annotations were exported in GFF3 format.

Gene prediction was conducted using Helixer v.0.3.6 with the land_plant model (Holst *et al*., 2026). Annotation completeness was evaluated using BUSCO with the eudicots_odb10 dataset (n = 2,326; Simão *et al*., 2015). Functional annotation of predicted genes was performed using BLASTx searches against the NCBI non-redundant (nr) protein database and domain annotation using InterProScan (Blum *et al*., 2021). Gene Ontology (GO) terms were assigned using Blast2GO (Conesa *et al*., 2005). Only high-confidence gene models were retained for downstream analyses.

Transposable elements (TEs) were identified using RepeatModeler2 (Flynn *et al*., 2020), which integrates RECON (Bao and Eddy, 2002), RepeatScout (Price *et al*., 2005), LTRharvest (Ellinghaus *et al*., 2008), and LTR_retriever (Ou and Jiang, 2018). A non-redundant, species-specific TE library generated by RepeatModeler2 was further classified using homology searches against the Dfam database (Storer *et al*., 2021), and TE annotation was performed using RepeatMasker v4.1.6. (Tarailo-Graovac and Chen, 2009).

### Variant detection and genome feature visualization

Illumina paired-end and mate-pair reads were aligned to the primary reference assembly using Bowtie2 with default parameters (Langmead and Salzberg, 2012). Alignments were coordinate-sorted, assigned read groups and checked for PCR duplicates using Picard (Broad Institute, 2019), and indexed with SAMtools (Danecek *et al*., 2021). Variants were called with GATK v.4.0.2.1 HaplotypeCaller (McKenna *et al*., 2010) using default parameters. VCF files were converted to BED format using BEDOPS (Neph *et al*., 2012). Genomic features were visualized using Circos (Krzywinski *et al*., 2009).

### Pollen nuclei isolation, 10x Genomics scRNA-seq library preparation and sequencing

The workflow followed the protocol described by Campoy *et al*. (2020), with minor modifications. Anthers were collected from fully mature flowers of the same individual used for genome sequencing and immersed in a buffer formulated for woody-species pollen (Loureiro *et al*., 2007) to release the pollen grains. Nuclei were obtained through a modified mechanical disruption approach: the pollen suspension was first passed through a 100 µm mesh for pre-filtration, and the grains were then ruptured by pressing them through a 30 µm mesh (Celltrics). The released nuclei were collected in pollen buffer and labeled with DAPI (4,6-diamidino-2-phenylindole; final concentration of 1 µg ml⁻¹). Sorting was carried out on a BD FACSAria Fusion flow cytometer fitted with a 70 µm nozzle, operating at a sheath pressure of 483 kPa. A total of 10,000 nuclei were collected into 23 µl of sheath fluid and subsequently loaded onto a 10x Chromium controller following the manufacturer’s protocol. Library construction followed the Chromium Single Cell 5′ workflow kit from 10x Genomics.

Sequencing was performed on an Illumina NovaSeq platform in 150 bp paired-end mode, yielding a total output of 100 Gb.

### Crossover identification

Crossover (CO) detection followed the pipeline outlined by Castellani *et al*. (2024). In brief, SNP markers across the genome were defined by aligning PacBio HiFi reads to haplotype 1 using minimap2 (Li, 2018), with SNPs subsequently called through ’bcftools mpileup’ and ’bcftools call’ (v.1.9; Danecek *et al*., 2021). Only allelic SNPs were retained as markers to discriminate between the two haplotypes; accordingly, variant-level information — including mapping quality, alternative-allele coverage and allele frequency — was compiled following conversion with SHOREmap (v.3.6) (Schneeberger *et al*., 2009), a tool that transforms SNP call files (.vcf) into a tab-delimited, more accessible text format.

To define genotyping markers in *S. tuberosa* pollen nuclei, scRNA-seq reads were first aligned to the haplotype 1 chromosome set using HISAT2 (v.2.1.0; Kim *et al*., 2019). For each nucleus, the paired reads were combined into a single-end FASTQ file, and alignment was run in single-end mode (-U), since the SNP-calling strategy fails to detect variants on reads whose mate pair remains unmapped. Prior to downstream analysis of the alignments, UMIs were extracted together with the cell barcodes; PCR duplicates were then removed by collapsing reads sharing identical UMIs, using the UMIcollapse tool (Liu, 2019).

SNP calling across all pollen nuclei followed the same procedure applied to the reference genome — namely, ’bcftools mpileup’ and ’bcftools call’ (v.1.9) — except that the ’--variants-only’ flag was omitted. Once SNPs had been called for each pollen nucleus, their positions along with reference- and alternative-allele counts were retrieved using ’bcftools query’. Cross-referencing the nucleus-level SNPs against the reference-defined markers yielded a set of reliable genotyping markers for that nucleus. CO identification was then carried out by adapting the haplotype-phasing approach described by Campoy *et al*. (2020).

### Orthology and synteny analysis

Orthologous gene clustering and functional analyses were performed using OrthoVenn3 (Sun *et al*., 2023). To investigate gene collinearity and infer syntenic relationships among Anacardiaceae genomes, we employed the GENESPACE v.1.2.3 pipeline (Lovell *et al*., 2022). Comparative analyses included four other Anacardiaceae species (*Anacardium occidentale*, *Pistacia vera*, *Pistacia integerrima* and *Mangifera indica*) and two representatives of Sapindaceae (*Acer negundo* and *Acer yangbiense*), used as outgroups. Genome assemblies of *M. indica*, *P. vera*, *P. integerrima*, *A. negundo* and *A. yangbiense* were downloaded from NCBI and annotated with Helixer, using the same procedure applied to *S. tuberosa*, to minimize differences arising from annotation methods. For *A. occidentale*, the v0.9 annotation available in Phytozome (Goodstein *et al*., 2012) was used, retaining only the longest transcript per gene (40,848 proteins).

An all-versus-all similarity search was performed using DIAMOND v.2.1.12 (Buchfink *et al*., 2021) with an e-value cutoff of 1 × 10⁻⁵ to identify homologous gene pairs across genomes. Syntenic blocks were then detected with MCScanX (Wang *et al*., 2012), which identifies collinear gene pairs based on both sequence similarity and conserved gene order. Syntenic orthogroups were inferred by integrating pairwise syntenic gene anchors across all genomes into a pan-genomic synteny graph.

To investigate whole-genome duplication (WGD) events, synonymous substitution rates (Ks) were estimated for collinear gene pairs within *S. tuberosa*, *M. indica* and *A. occidentale* (paralogs), and between *M. indica* and *A. occidentale* and between *S. tuberosa* and *M. indica* (orthologs). For each species, the longest coding sequence (CDS) per gene was retained and translated. All-against-all protein similarity searches were performed with DIAMOND v.2.1.12 (sensitive mode; e-value ≤ 1 × 10^−5^; up to five hits per query; Buchfink *et al*., 2021), and collinear blocks of at least five gene pairs were identified with MCScanX using default parameters (Wang *et al*., 2012). For each collinear gene pair, protein sequences were aligned with MUSCLE v.5.3 (Edgar, 2022) and back-translated into codon alignments; codons with gaps were removed, and pairs with fewer than 150 aligned nucleotides were discarded. Ks was estimated with the Yang and Nielsen (2000) method implemented in yn00 (PAML; Yang, 2007). Ks distributions (0.01 < Ks < 2) were summarized by Gaussian kernel density estimation (bandwidth = 0.03), and 95% confidence intervals of the main peaks were obtained from 100 bootstrap replicates of gene pairs.

To estimate divergence times, we performed a phylogenomic analysis based on the nuclear genomes of the seven species. Single-copy orthologous genes shared by all species were aligned with MAFFT v.7.222 on an online platform (Katoh *et al*., 2019). Phylogenetic relationships were inferred by maximum likelihood (ML) under a General Time Reversible + Gamma + Invariant Sites (GTR + G + I) substitution model (Abadi *et al*., 2019). The ML analysis was performed using RAxML-HPC2 (Stamatakis, 2014) on the CIPRES Science Gateway (Miller *et al*., 2010). Divergence times were estimated using BEAST v.1.8.3 (Drummond *et al*., 2012), with the ML tree used as a fixed topology. Calibration was based on the Anacardiaceae/Sapindaceae divergence (76.0 Ma; Li *et al*., 2019). An uncorrelated relaxed lognormal clock (Drummond and Rambaut, 2007) and a birth–death speciation model (Gernhard, 2008) were applied. Two independent runs of 100,000,000 generations were performed, sampling every 10,000 generations. After discarding 25% of samples as burn-in, the independent runs were combined, and a maximum clade credibility tree was constructed using TreeAnnotator v.1.8.3 (Rambaut and Drummond, 2013). Convergence of independent chains and adequate sampling of all parameters were assessed in Tracer v.1.7 (Rambaut *et al*., 2018), with effective sample sizes (ESS) ≥ 200 considered sufficient.

### Gene family expansion and contraction analysis

Gene family expansion and contraction across the Anacardiaceae phylogeny were inferred using the integrated evolutionary analysis module of OrthoVenn3 (Sun *et al*., 2023). A phylogenetic tree was reconstructed within the platform using FastTree2 (Price *et al*., 2010), based on single-copy orthologous clusters identified across the seven species. The tree was time-calibrated in OrthoVenn3 using node ages from the nuclear phylogenomic BEAST analysis (Fig. 5) for the root (Anacardiaceae/Sapindaceae divergence), the split between *S. tuberosa* and the remaining Anacardiaceae, and the *M. indica*–*A. occidentale* split. Gene family expansion and contraction were then estimated using CAFE5 (Mendes *et al*., 2020), embedded in the OrthoVenn3 pipeline, based on these divergence time estimates, with significance assessed at a p-value threshold of 0.05. Gene families showing significant expansion in the *S. tuberosa* lineage were subsequently retrieved and functionally annotated using BLASTx searches against the NCBI nr database, InterProScan (Blum *et al*., 2021), and Blast2GO (Conesa *et al*., 2005). Expanded genes located within five genes of another member of the same family on the same chromosome were considered tandem duplicates.

## RESULTS

### Genome sequencing and assembly

A total of 1.8 million high-fidelity PacBio HiFi reads (Supplementary Table S1) were used for genome assembly, resulting in a total assembly size of 336 Mb, of which 288 Mb were anchored onto the 16 chromosomes. The Hi-C contact map revealed a strong interaction signal along the main diagonal, supporting a high-quality chromosome-scale assembly with 16 well-defined chromosomes (Fig. 1A). The clear compartmentalization of each chromosome indicates accurate scaffolding and correct ordering and orientation of contigs. Interchromosomal signals were weak overall and concentrated in discrete X-shaped foci connecting the putative centromeric regions of different chromosomes, a pattern consistent with the clustering of centromeres within the interphase nucleus that is commonly observed in plant Hi-C maps, rather than with scaffolding errors. The assembly generated primary and alternate haplotypes, and the primary assembly was used for downstream analyses. Genome completeness assessed using BUSCO showed high recovery of conserved orthologs (Fig. 1B): 2,298 of the 2,326 BUSCO groups (eudicots_odb10) were identified as complete in the genome assembly (98.8%), comprising 2,286 single-copy and 12 duplicated genes, with only 6 fragmented and 22 missing. Similarly, the predicted gene set exhibited 2,301 complete BUSCOs (98.9%), comprising 2,283 single-copy and 18 duplicated genes, with 16 fragmented and 9 missing.

**Fig. 1.**
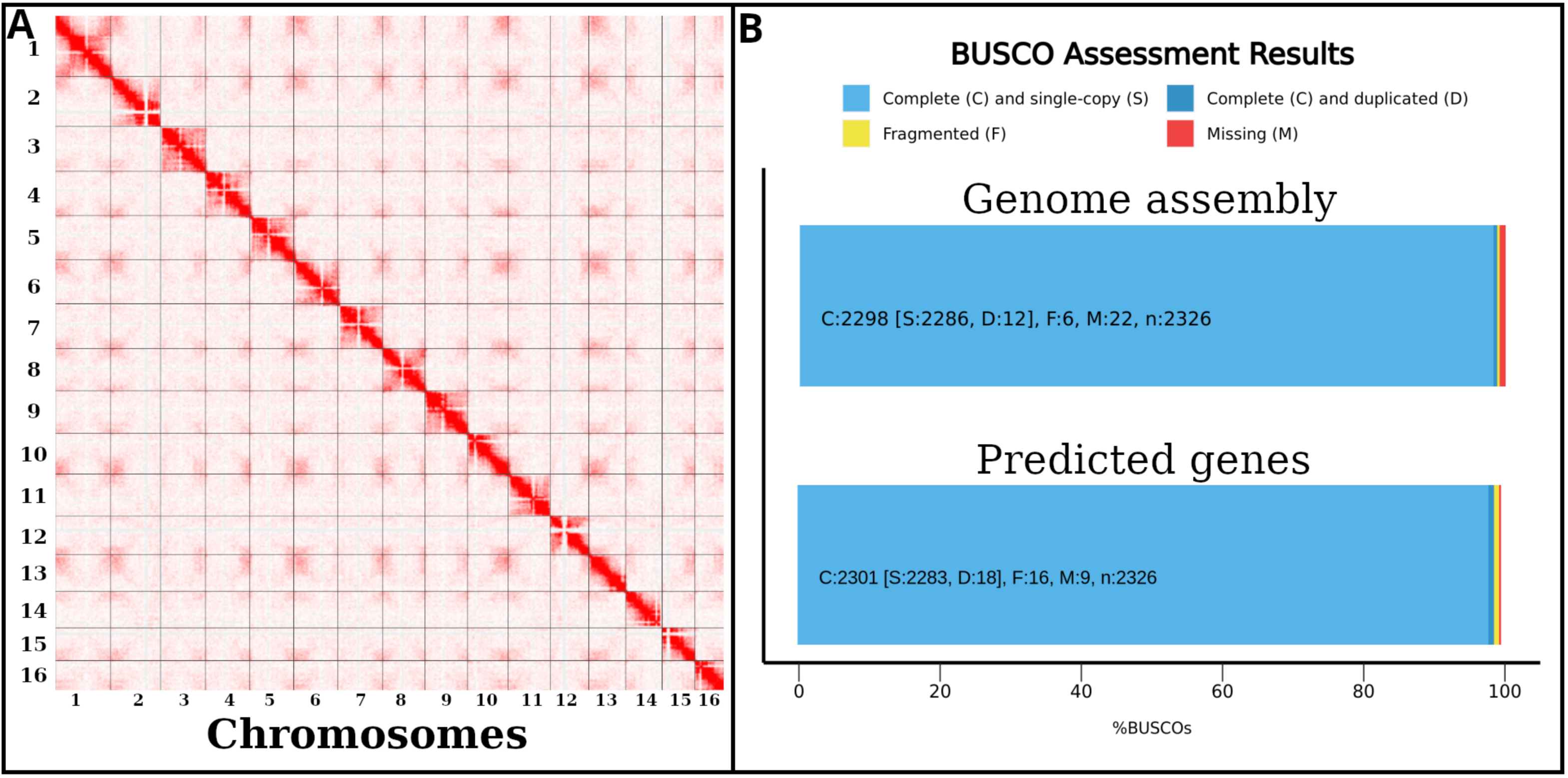
Genome assembly quality and completeness assessment. (A) Genome-wide contact map (Hi-C) showing chromosomal interactions for the assembled genome. (B) BUSCO assessment of genome completeness for the assembly and predicted gene set. Alt text: Two-panel figure. Panel A is a Hi-C contact heat map with 16 bright squares along the diagonal, one per chromosome. Panel B is a bar chart of BUSCO results showing nearly all conserved genes complete in both the assembly and the predicted gene set.

Chromosome sizes ranged from 27.3 to 12.7 Mb, totaling 288,380,347 bp. The two haplotypes resolved by the assembly reflect the high heterozygosity of the sequenced individual. Single-nucleotide polymorphisms (SNPs) were identified by mapping short reads onto the primary reference genome using GATK, yielding a total of 1,780,712 heterozygous SNPs, corresponding to a heterozygosity rate of 0.62%, distributed across all chromosomes.

The Circos plot provides an integrated view of genomic features across the 16 chromosomes (Fig. 2). Satellite DNA occurs in discrete blocks across most chromosomes, with pronounced enrichment in specific regions of certain chromosomes (e.g., chr2 and chr12), while other chromosomes display comparatively reduced signal, whereas SNP density remains relatively uniform along the genome, with only modest local fluctuations. Predicted genes are broadly distributed across chromosomes, with variation in local density. Transposable elements show distinct patterns, with Ty1/copia and Ty3/gypsy elements widely dispersed and enriched in specific regions, while LINE elements are less abundant. GC content remains relatively stable across chromosomes, with moderate local variation. Inner links indicate intra-genomic synteny, suggesting conserved genomic regions and segmental duplications.

**Fig. 2.**
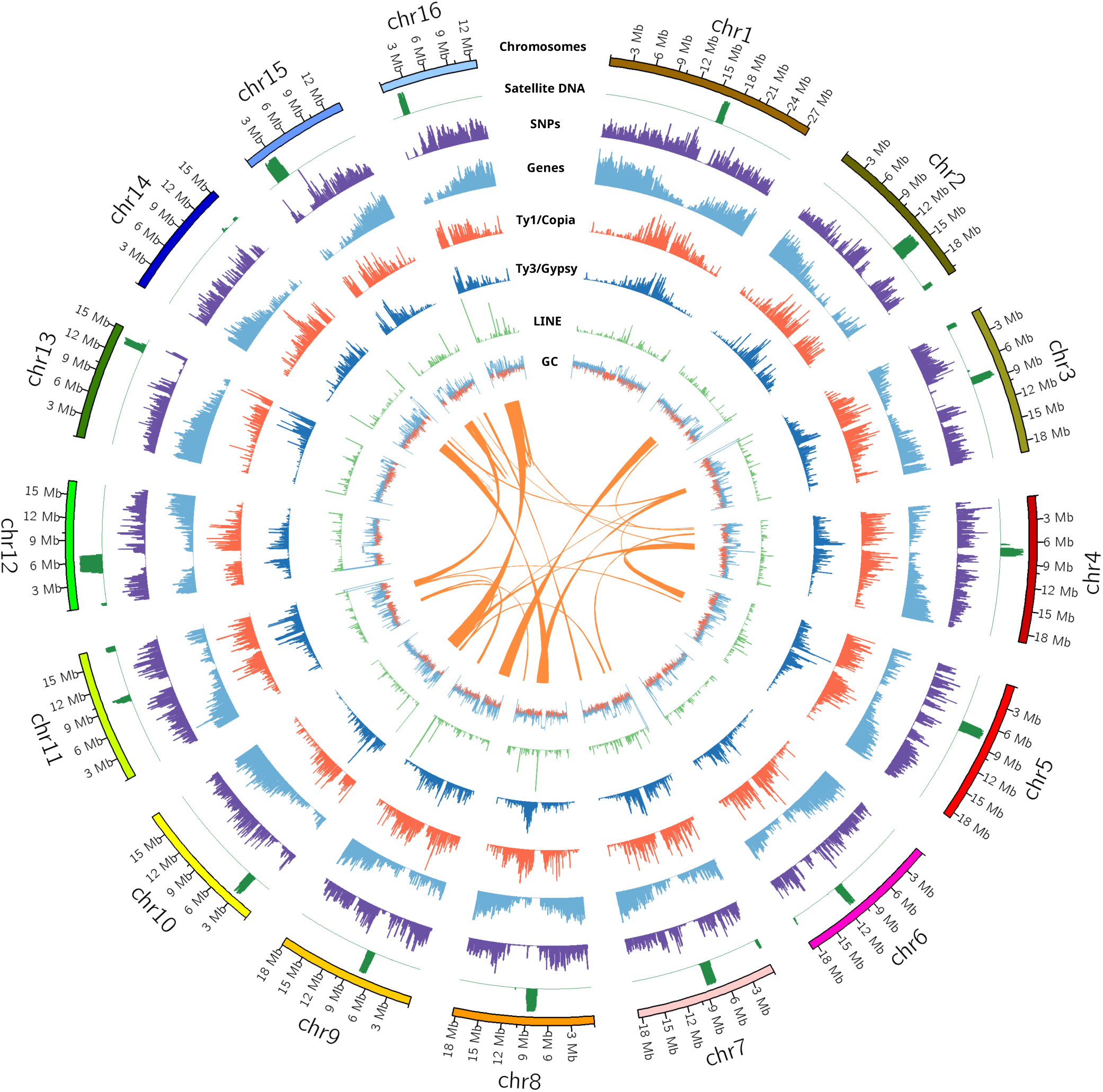
Circos representation of genome features and structural organization. Links in the center represent intra-genomic synteny (self-synteny) relationships across chromosomes. Alt text: Circular Circos plot of the 16 chromosomes with concentric tracks for satellite DNA, SNP density, gene density, transposable elements and GC content, and internal links connecting syntenic regions within the genome.

### Genome annotation

Repetitive DNA was identified using RepeatModeler2 and classified through homology-based searches against repeat libraries using RepeatMasker. Repetitive sequences accounted for 52.01% of the genome assembly, including 25.94% de novo repeat families, 9.11% Ty3/gypsy, 6.95% satellite DNA, 6.43% Ty1/copia, 3.29% DNA transposons, and 0.29% LINEs (Supplementary Table S2). Gene prediction identified 24,232 protein-coding genes, occupying 33.40% of the genome, as well as multiple copies of 35S ribosomal DNA on chromosome 14 and 5S ribosomal DNA on chromosome 4. Additionally, 494 transfer RNA (tRNA) genes were annotated.

The distribution of SNPs across annotation categories revealed that the highest number of variants is associated with gene regions and unknown repeat sequences (Fig. 3A). In contrast, transposable element classes such as Ty1/copia, Ty3/gypsy, and LINEs, as well as SSRs, satellites, and simple repeats, contribute relatively fewer SNPs (Fig. 3A). At the chromosomal level, SNPs are broadly distributed, with no strong chromosome-specific bias (Fig. 3A).

**Fig. 3.**
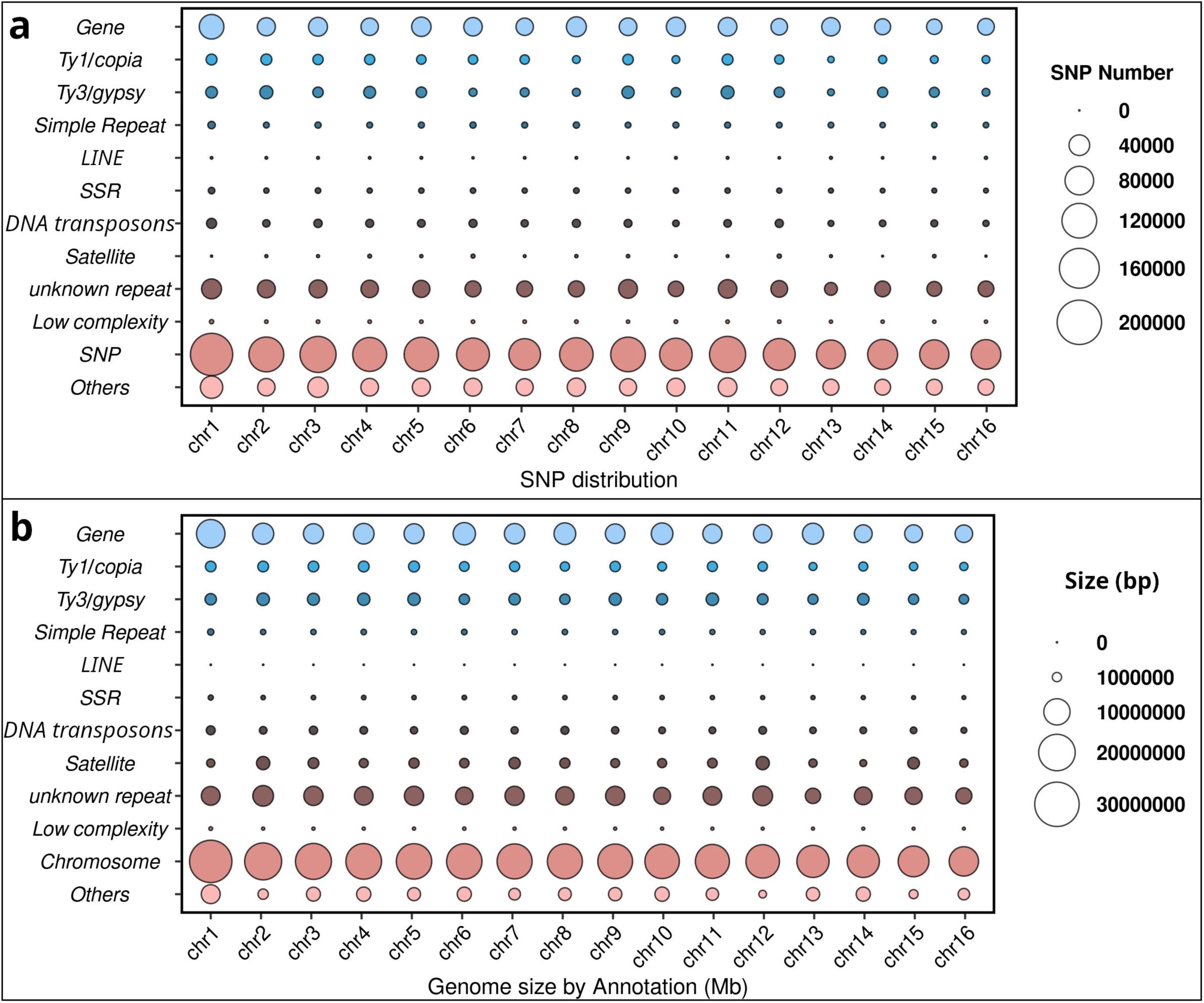
Distribution of SNPs and genomic content across annotation classes and chromosomes of *Spondias tuberosa*. (A) Number of SNPs in each annotation class per chromosome (chr1– chr16); circle size is proportional to the number of SNPs. (B) Genomic span of each annotation class per chromosome; circle size is proportional to the size of each class. In (A) and (B), “SNP” and “Chromosome” indicate the total number of SNPs and the total size of each chromosome, respectively. Genome-wide values for each class are given in Supplementary Table S2. Alt text: Two bubble charts with annotation classes in columns and chromosomes 1 to 16 in rows; bubble size shows SNP number in panel A and genomic span in panel B, with gene regions and unknown repeats having the largest bubbles.

Regarding genome composition, the largest fractions of the genome are represented by gene regions (33.40%), unknown repeats (25.94%) and unclassified regions (13.48%), whereas specific repeat families occupy smaller proportions (Table S2). The chromosomal distribution of genomic size by annotation confirms a relatively uniform organization across chromosomes, with major contributions from gene regions, unknown repeats and unclassified regions (Fig. 3B).

### CO landscape

Quantification of meiotic crossover (CO) events across the 16 chromosomes identified 1,740 COs from 153 pollen nuclei, corresponding to a mean of 11.37 COs per pollen nucleus — exceeding the ∼8.0 COs per nucleus expected under a one-CO-per-bivalent assurance model. COs were detected across all chromosomes but were unevenly distributed along their physical length, resulting in pronounced chromosome- and region-specific variation in recombination rate (Supplementary Fig. S1). The genome-wide recombination rate was estimated at 4.01 cM Mb⁻¹, corresponding to 1,137 cM across the 284 Mb of haplotype 1, used as the reference for crossover detection. Recombination was generally enriched in interstitial and distal chromosome regions, while markedly reduced in pericentromeric regions, indicating that meiotic recombination is highly heterogeneous across the *Spondias* genome. Notably, regions of reduced CO frequency coincided with regions enriched in repetitive DNA, particularly transposable elements and satellite DNA (Supplementary Fig. S2), suggesting an association between chromosome repetitive architecture and the spatial organization of meiotic recombination.

### Comparative analysis of orthologous gene clusters and functional annotation

Orthologous cluster analysis using OrthoVenn3 across Anacardiaceae species revealed both shared and species-specific gene families (Fig. 4A, B). A substantial number of clusters were conserved across all species, representing core gene functions, while unique clusters were identified for each species, indicating lineage-specific gene content. The number of orthologous clusters varied among species, with *Pistacia vera* and *Pistacia integerrima* showing the highest numbers, whereas *Spondias tuberosa* exhibited the fewest (Fig. 4B).

**Fig. 4.**
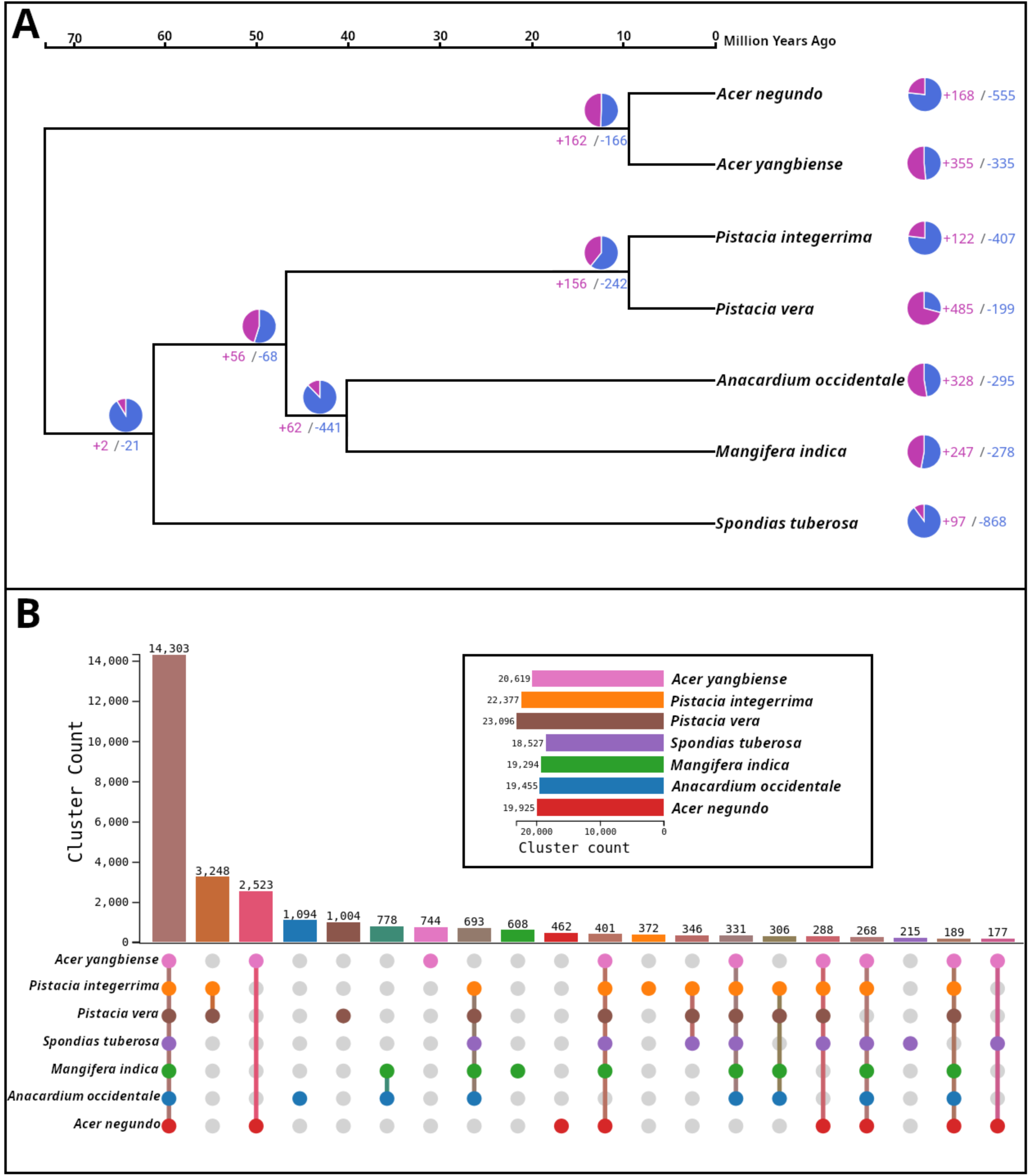
Orthologous gene clustering and gene family evolution in *Spondias tuberosa* and six other Sapindales species. (A) Phylogenetic tree reconstructed in OrthoVenn3, with the numbers of expanded (+) and contracted (−) gene families inferred by CAFE5 for each branch. (B) Distribution of orthogroups across species. The bar plot shows the total number of gene clusters per species, and the UpSet plot illustrates shared and species-specific orthogroups. Divergence times in (A) were constrained using node ages from the analysis shown in Fig. 5. Alt text: Panel A is a phylogenetic tree of seven Sapindales species with numbers of expanded and contracted gene families on each branch. Panel B is an UpSet plot showing gene clusters shared among species and unique to each species, with a bar chart of total clusters per species.

**Fig. 5.**
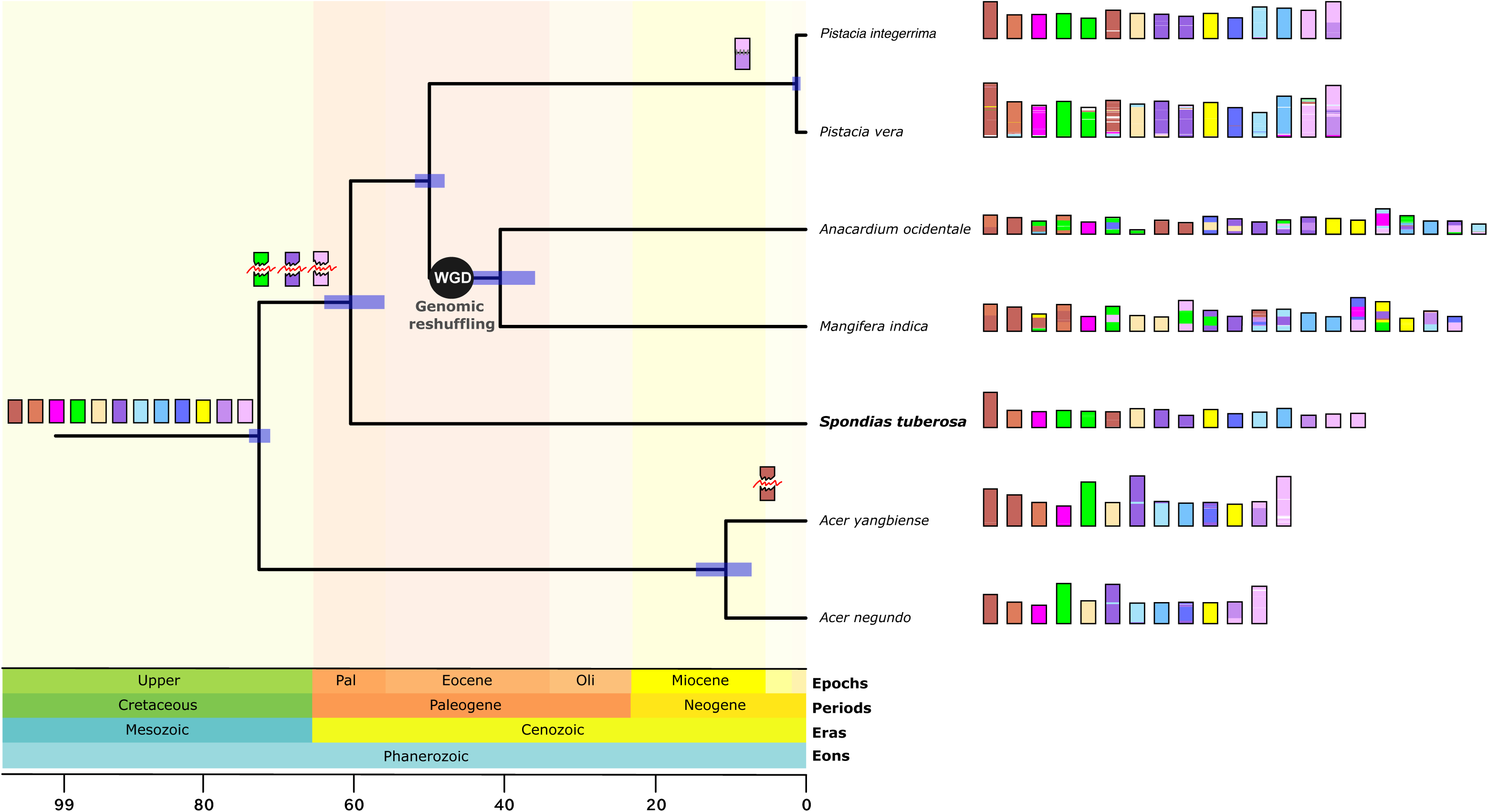
Phylogenetic relationships, divergence time estimates, and genome evolution of *Spondias tuberosa* and related species. The tree is time-calibrated with geological periods indicated in the background. Divergence times were estimated with BEAST from nuclear genome data and are shown in million years (Ma); blue bars represent 95% highest posterior density (HPD) intervals of node ages, and the “WGD” node indicates a shared whole-genome duplication event followed by genomic reshuffling. Chromosome diagrams illustrate genome organization and lineage-specific rearrangements. Alt text: Time-calibrated phylogenetic tree of Spondias tuberosa and six related species with a geological time scale, node age bars, a marked whole-genome duplication on the Mangifera– Anacardium branch, and chromosome diagrams beside each species.

Gene Ontology (GO) annotation revealed that most genes were associated with biological processes, followed by cellular components and molecular functions (Supplementary Fig. S3).

Within biological processes, cellular and metabolic processes were predominant, while binding and catalytic activity were the main molecular functions. The 97 gene families significantly expanded in *S. tuberosa* comprised 448 genes (Supplementary Table S3), most of which (369 genes; 82%) were located within five genes of another member of the same family, indicating that expansion occurred predominantly through tandem duplication. These families included genes associated with environmental response, signaling, and development. Expanded genes included receptor-like kinases (RLKs), disease resistance proteins, gibberellin 20-oxidases, F-box proteins (including S-locus F-box proteins), glutathione *S*-transferases, and genes involved in cell wall organization (cellulose synthase-like proteins), transport and secondary metabolism (terpene synthases and cytochrome P450 monooxygenases), suggesting that gene family expansion in this species may be linked to stress response, regulatory processes, and metabolic adaptation. In contrast, 868 gene families were contracted in *S. tuberosa*, 605 of which contained no *S. tuberosa* genes.

### Genome evolution and phylogenetic relationships

Phylogenetic analysis places *Spondias tuberosa* as sister to the clade comprising *Pistacia* and the *Mangifera–Anacardium* lineage, consistent with the placement of *Spondias* in subfamily Spondioideae, sister to Anacardioideae, while *Acer* species — representing a distinct family (Sapindaceae) — occupy a more distant, outgroup position (Fig. 5). The divergence between Anacardiaceae and Sapindaceae was dated to the Late Cretaceous (c. 73 Ma), the split between *S. tuberosa* and the remaining Anacardiaceae to the Paleocene (c. 60 Ma), the split between *Pistacia* and the *Mangifera–Anacardium* lineage to the early Eocene (c. 50 Ma), and the divergence between *Mangifera* and *Anacardium* to the middle Eocene (c. 41 Ma), whereas the two *Acer* species diverged in the Miocene (c. 11 Ma) (Fig. 5). Ks distributions of collinear paralogs showed a recent peak in *M. indica* (Ks = 0.26) and *A. occidentale* (Ks = 0.30), indicative of a whole-genome duplication (WGD), which was absent in *S. tuberosa*, whose paralogs retained only the older peak (Ks ≈ 1.5) attributed to the γ triplication shared by core eudicots (Fig. 6A). Orthologs between *M. indica* and *A. occidentale* peaked at a lower Ks (0.18) than their paralogs, indicating that the WGD preceded the divergence of these two genera, whereas *S. tuberosa*–*M. indica* orthologs peaked at a higher Ks (0.37), placing the WGD after the split between the *Spondias* and mango–cashew lineages. Together, these results indicate a WGD shared by *Mangifera* and *Anacardium*, followed by extensive genomic reshuffling (Fig. 5). Chromosomal comparisons revealed an overall conserved genome structure across Anacardiaceae, with lineage-specific rearrangements. In *S. tuberosa*, chromosome structure is largely conserved relative to related species, with only localized structural variation.

**Fig. 6.**
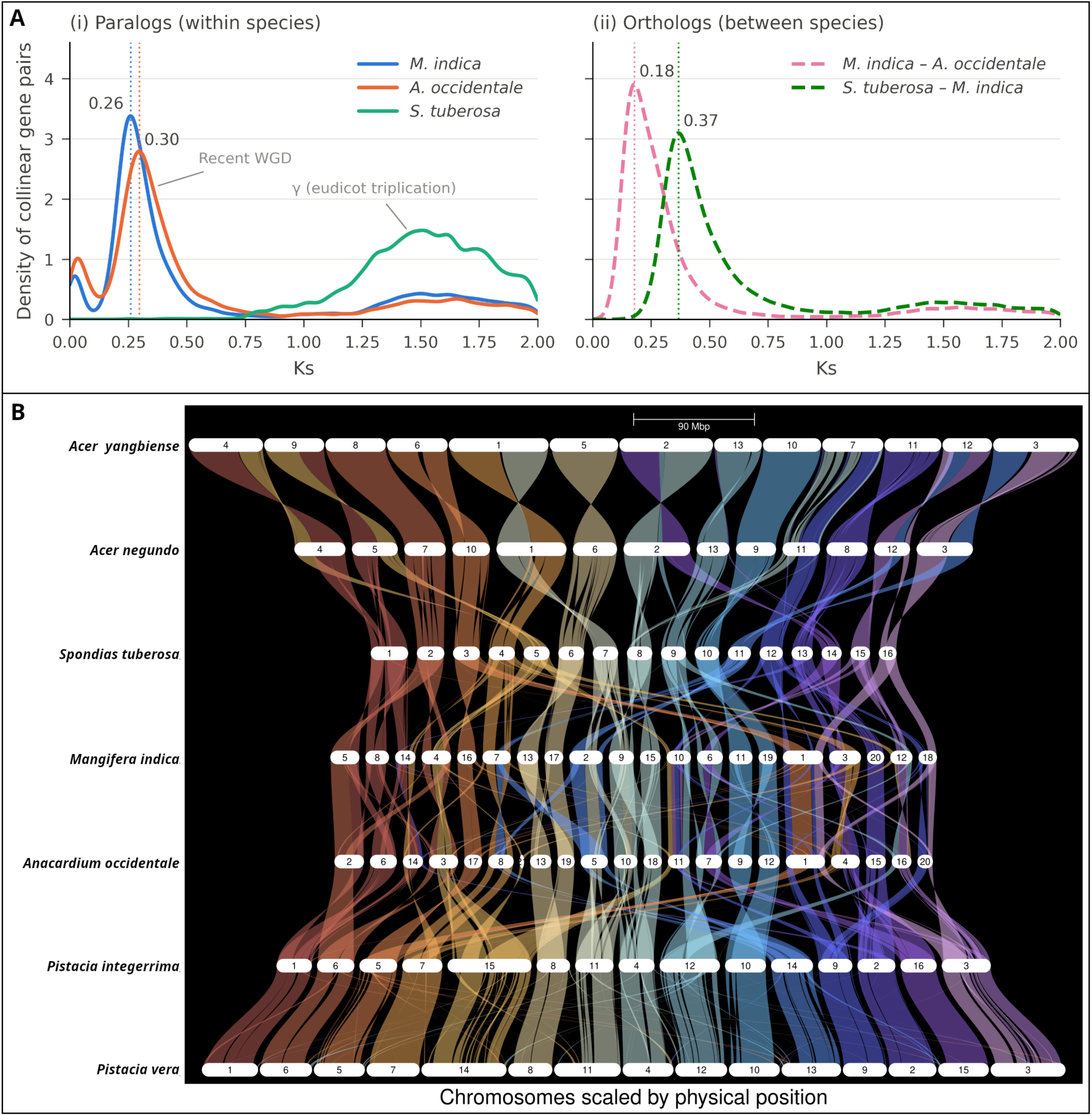
Synonymous substitution and synteny analyses of *Spondias tuberosa* and related species. (A) Distribution of synonymous substitution rates (Ks) for collinear gene pairs: (i) paralogs and (ii) orthologs. (B) Synteny relationships among species. Chromosomes are shown as horizontal bars, and colored ribbons represent conserved syntenic blocks. Alt text: Panel A shows density curves of synonymous substitution rates for paralogous and orthologous gene pairs, with a recent peak in mango and cashew absent in Spondias tuberosa. Panel B shows horizontal chromosome bars for each species connected by coloured ribbons of conserved synteny.

Synteny analysis revealed extensive collinearity among species (Fig. 6B). High levels of synteny were observed between closely related species, particularly between *Pistacia vera* and *Pistacia integerrima*, as well as between *S. tuberosa* and *M. indica*. In contrast, *Acer* species showed fewer and more fragmented syntenic blocks. Despite this overall conservation, several chromosomal rearrangements were detected, including translocations and segment reorganization, indicating that genome evolution in Anacardiaceae has been shaped jointly by whole-genome duplication, conservation of ancestral syntenic blocks, and lineage-specific structural modifications.

## DISCUSSION

### Spondias tuberosa genome

The current assembly (336 Mb total; 288 Mb anchored) is smaller than the genome size previously estimated for *S. tuberosa* by flow cytometry (513.01 Mb; Martins *et al*., 2025), though closer to the k-mer-based estimates reported in the same study (349–360 Mb). This gap is likely explained, at least in part, by the substantial satellite DNA content previously characterized in the species, which reached 7.10% of the genome using a genome-skimming approach (Martins *et al*., 2025) — a proportion consistent with the 6.95% satellite DNA identified in the present chromosome-scale assembly (Table S2). Satellite monomers are typically homogenized through concerted evolution, resulting in low sequence divergence among repeat copies; in agreement with this, satellite regions harboured very few SNPs relative to their genomic span (Fig. 3). This limited intra-array sequence variation constrains the ability of long-read assemblers, which rely on sequence divergence to distinguish and correctly phase repeat copies, likely resulting in the collapse or under-representation of large satellite arrays in the final assembly — a plausible explanation for part of the discrepancy between the assembled genome size and previous flow cytometry-based estimates. Because satellite DNA alone (c. 7% of the genome) cannot account for the full difference of c. 175 Mb, the collapse of other high-copy repeats, such as LTR retrotransposons, probably also contributes.

The chromosome-scale assembly revealed 16 well-defined chromosomes, consistent with the conserved karyotypic organization described for species within the genus (Almeida *et al*., 2007). The high BUSCO completeness (>98%) further supports the quality of the assembly and is comparable to other high-quality genomes reported for the family (Zeng *et al*., 2019; Wang *et al*., 2020). The heterozygosity rate of 0.62% indicates a high level of heterozygosity in the sequenced individual, consistent with the outcrossing, self-incompatible mating system of *S. tuberosa* (Nadia *et al*., 2007).

The distribution of genomic features along chromosomes shows patterns consistent with other Anacardiaceae genomes. Transposable elements, particularly Ty1/copia and Ty3/gypsy, are widely distributed and represent a major component of genome structure, as reported in *M. indica* and *P. vera* (Zeng *et al*., 2019; Wang *et al*., 2020). Satellite DNA forms one major block on most chromosomes (Fig. 2), which likely corresponds to centromeric and pericentromeric regions, where it may contribute to chromosomal stability.

The intra-genomic synteny detected in *S. tuberosa* (Fig. 2) reflects conserved duplicated blocks and segmental duplications. Because *S. tuberosa* lacks the recent WGD detected in mango and cashew (Fig. 6A), these blocks most likely represent remnants of the ancient γ triplication shared by core eudicots, together with more recent segmental duplications, rather than a lineage-specific polyploidy event.

### Whole-genome duplication and genome architecture in Anacardiaceae

Our Ks analyses indicate that the mango–cashew lineage experienced a WGD that is absent from *S. tuberosa*. The recent paralog peaks of *M. indica* (Ks = 0.26) and *A. occidentale* (Ks = 0.30) are older than the divergence of the two genera (ortholog Ks = 0.18) but younger than their divergence from *Spondias* (ortholog Ks = 0.37), placing the event between c. 60 and c. 41 Ma according to our dated phylogeny (Fig. 5). This is consistent with the recent WGD reported for mango (Wang *et al*., 2020) and with the higher chromosome numbers of *Mangifera* (n = 20) and *Anacardium* (n = 21) relative to *Spondias* (n = 16) and *Pistacia* (n = 15). Bai *et al*. (2022) proposed that *Toxicodendron vernicifluum* shares a WGD with mango, whereas no lineage-specific WGD was detected in pistachio. Because *Toxicodendron* is more closely related to *Pistacia* than to *Mangifera* (Bai *et al*., 2022), the timing of the event relative to the divergence of *Pistacia* remains unresolved. Including *Pistacia* and *Toxicodendron* paralogs in future Ks analyses will be required to determine whether the WGD predates this split and was followed by extensive fractionation in some lineages, or occurred only in the *Mangifera–Anacardium* lineage, as depicted in Fig. 5.

The absence of a recent WGD in *S. tuberosa* supports our hypothesis that its genome retains a comparatively conserved architecture. Its chromosomes show largely one-to-one syntenic relationships with those of the *Acer* outgroups (Fig. 6B), and only 12 of the 2,326 BUSCO genes (0.5%) were duplicated in the assembly (Fig. 1B). In the absence of polyploidy, gene family expansion in *S. tuberosa* proceeded mainly through tandem duplication: 82% of the genes in expanded families were located near another member of the same family, as in the gibberellin 20-oxidase clusters on chromosomes 7 and 14 and the subtilisin-like protease cluster on chromosome 16 (Table S3). Tandem duplication is a major source of lineage-specific expansion of stress-responsive gene families in plants (Hanada *et al*., 2008). Together with its position as sister to the remaining Anacardiaceae sampled, this unduplicated and largely collinear architecture makes *S. tuberosa* a suitable reference for reconstructing ancestral genome organization and for transferring genomic information across the family.

### Crossover landscape in a perennial tree

In perennial trees, characterizing recombination is particularly challenging, as conventional linkage mapping relies on segregating populations that are time-consuming to establish in species with long juvenile periods. The pollen-based landscape obtained here showed recombination markedly reduced in pericentromeric regions and elevated in interstitial regions, a pattern typical of monocentric chromosomes, with a genome-wide rate of 4.01 cM Mb⁻¹.

This rate is higher than the values of approximately 1.5–3.0 cM Mb⁻¹ reported for other woody perennials, including apple, peach, cacao, grapevine and *Eucalyptus* (Tiley and Burleigh, 2015). Two factors should be considered when interpreting this difference. First, our estimate is derived exclusively from male meiosis, and recombination rates frequently differ between sexes (Lenormand and Dutheil, 2005); second, recombination rate per physical distance tends to decrease with increasing genome size (Tiley and Burleigh, 2015), so higher values are expected in the comparatively compact genome of *S. tuberosa*. This pollen-based approach nevertheless allows recombination landscapes to be characterized in *Spondias* without establishing a segregating population and, together with the genome assembly, provides a resource for future genetic mapping and breeding.

### Expansion of gene families in *S. tuberosa*

Several of the families expanded in *S. tuberosa* are related to environmental perception and stress tolerance. Receptor-like kinases (RLKs) — especially leucine-rich repeat and serine/threonine kinase types — are among the expanded families, suggesting a heightened capacity for sensing environmental signals and transducing them into downstream responses. These receptors are central to recognizing extracellular stimuli and triggering signaling cascades associated with both pathogen recognition (Boutrot and Zipfel, 2017) and abiotic stress, including drought and osmotic imbalance (Osakabe *et al*., 2013; Gong *et al*., 2020).

Gene families associated with detoxification and oxidative stress mitigation were also expanded, including five glutathione *S*-transferases (GSTs). These enzymes contribute to redox homeostasis by neutralizing reactive oxygen species (ROS) that accumulate under drought conditions (Gill and Tuteja, 2010), and their expansion suggests a strengthened capacity to counteract oxidative damage — a feature commonly associated with drought-tolerant genotypes (Nianiou-Obeidat *et al*., 2017; Hasanuzzaman *et al*., 2020).

Hormonal pathways emerge as another adaptive axis. Gibberellin 20-oxidases, which catalyse key steps in the biosynthesis of bioactive gibberellins, formed one of the largest expanded families (21 genes), suggesting fine-tuning of growth regulation, and two genes encoding PYL-type abscisic acid (ABA) receptors (PYL11- and PYL12-like) were also expanded. ABA signaling is a central regulator of drought responses, governing processes such as stomatal closure and the induction of stress-responsive genes (Cutler *et al*., 2010; Yoshida *et al*., 2014), thereby allowing the plant to balance growth with survival under limited water availability.

Cell wall remodeling genes — including xyloglucan endotransglucosylase/hydrolases and cellulose synthase-like proteins — were expanded as well, indicating structural adjustments that help preserve cell wall integrity and flexibility under dehydration (Tenhaken, 2015). The expansion of secondary metabolism genes, such as terpene synthases (e.g. nerolidol and caryophyllene synthases), cytochrome P450 monooxygenases and UDP-glycosyltransferases, further points to an enhanced capacity to synthesize protective compounds with antioxidant and ecological functions (Vogt, 2010; Tholl, 2015). Subtilisin-like proteases (16 genes), which participate in development, cell wall modification and immunity (Schaller *et al*., 2018), and germin-like proteins (12 genes) were also among the largest expanded families.

Defense-related gene families — including disease resistance proteins (33 genes) and thaumatin-like proteins — showed a similar pattern of expansion, consistent with the recognized overlap between biotic and abiotic stress signaling networks (Fujita *et al*., 2006). Lastly, the expansion of ERF transcription factors and of ubiquitin-related regulators, notably F-box proteins (21 genes), may increase regulatory versatility in responses to environmental challenges. Eight of these F-box genes encode S-locus F-box proteins, which in several eudicot lineages participate in S-RNase-based gametophytic self-incompatibility (Kubo *et al*., 2010). Notably, a family of eight T2 ribonucleases — the protein family to which S-RNases belong — was also expanded, and several of these genes are interleaved with S-locus F-box genes on chromosomes 3 and 14 (Table S3), an arrangement reminiscent of S-RNase-based S-loci. These findings are noteworthy given the self-incompatibility reported for *S. tuberosa* (Nadia *et al*., 2007), although the role of these genes in *Spondias* remains to be tested.

Overall, these expansions should be interpreted with caution: the *S. tuberosa* terminal branch showed the fewest expansions among the seven species (Fig. 4A), and several of the largest expanded families are primarily associated with biotic interactions; the expansions identified here should therefore be viewed as a targeted set of candidate genes rather than as evidence of a genome-wide adaptive signature. Some of these expansions may contribute to the species’ resilience to the recurrent drought episodes characteristic of the Caatinga biome, but functional validation — for example, expression profiling under water deficit — will be required to test these associations. Because all species except *A. occidentale* were annotated with the same Helixer pipeline, differences in annotation methods are unlikely to account for most of the inferred changes in gene family size; nevertheless, the absence of *S. tuberosa* genes in 605 of the 868 contracted families suggests that part of these contractions may reflect annotation or assembly differences rather than true gene loss.

## CONCLUSIONS

The *Spondias tuberosa* genome lacks the recent WGD shared by mango and cashew and retains a largely collinear architecture; recombination is concentrated in interstitial chromosome regions, whereas pericentromeric regions are enriched in transposable elements and satellite DNA. Gene family expansion, driven mainly by tandem duplication, identified candidate genes related to environmental perception, defense and self-incompatibility, providing a foundation for functional studies and for comparative and applied genomics in Anacardiaceae.

## SUPPLEMENTARY DATA

Supplementary data are available at *Annals of Botany* online and consist of the following. Table S1: summary of sequencing data used for genome assembly. Table S2: genome composition across chromosomes in *Spondias tuberosa*. Table S3: functional annotation of the 448 genes in the 97 gene families expanded in *S. tuberosa*. Figure S1: crossover (CO) landscape along the 16 chromosomes of *S. tuberosa* (haplotype 1), inferred from 153 pollen nuclei. Figure S2: Circos representation of genome features, including crossover distribution, for haplotype 1 of *S. tuberosa*. Figure S3: Gene Ontology (GO) annotation of the predicted protein-coding genes of *S. tuberosa*, showing the distribution of functional categories in Biological Process, Molecular Function and Cellular Component.

## AUTHOR CONTRIBUTIONS

C.A. and A.M. designed the research. E.L., D.S., G.R., J.V.S., A.B. and A.M. performed the experiments. All authors contributed to data analysis and interpretation. C.A. wrote the manuscript with input from all co-authors.

## DATA AVAILABILITY

The genome assembly, gene annotation and raw sequencing data generated in this study (PacBio HiFi, Omni-C, Illumina and pollen single-nucleus RNA-seq reads) are available at NCBI under BioProject PRJNA935689.

## ACKNOWLEDGEMENTS

We thank the Federal University of Alagoas for laboratory facilities and scientific support.

