## Supplementary tables for "The chromosome-scale genome of the drought-adapted Caatinga tree *Spondias tuberosa* reveals a conserved, unduplicated architecture within Anacardiaceae"

**Contents**

**Table S1.** Summary of sequencing data used for genome assembly.

**Table S2.** Genome composition across chromosomes in *Spondias tuberosa*.

**Table S3.** Functional annotation of the 448 genes in the 97 gene families expanded in *S. tuberosa*.

**Figure S1.** Crossover (CO) landscape along the 16 chromosomes of *S. tuberosa* (haplotype 1), inferred from 153 pollen nuclei.

**Figure S2.** Circos representation of genome features, including crossover distribution, for haplotype 1 of *S. tuberosa*.

**Figure S3.** Gene Ontology (GO) annotation of the predicted protein-coding genes of *S. tuberosa*.

Table S1. Summary of sequencing data and coverage for genome assembly

| **Metric** | **Illumina (short reads)** | **PacBio HiFi (long reads)** |
|---|---|---|
| Number of reads | 93,315,956 | 1,800,438 |
| Paired-end reads | 28,132,784 | – |
| Mate-pair reads (1–12 kb) | 65,184,172 | – |
| Average read length | 100 bp | 16.3 kb (102–50,031 bp) |
| Total bases | 18,7 Gb | 29.4 Gb |

Table S2. Genome composition across chromosomes in *Spondias tuberosa*.

| Chromosome | Size (bp) | Gene regions | Ty1/copia | Ty3/gypsy | DNA transposons | LINE | Satellite DNA | De novo family | Low complexity | SSRs | Other regions |
|---|---|---|---|---|---|---|---|---|---|---|---|
| chr1 | 27,344,340 | 11,912,402 | 1,401,854 | 1,644,499 | 937,349 | 51311 | 779,571 | 4,941,944 | 88,992 | 237,139 | 4,989,342 |
| chr2 | 20,609,735 | 6,341,074 | 1,434,873 | 2,043,416 | 637,489 | 63771 | 2,332,690 | 6,048,211 | 52,430 | 154,594 | 1,298,330 |
| chr3 | 19,473,299 | 5,638,287 | 1,441,504 | 1,866,558 | 797,973 | 39497 | 1,459,729 | 5,211,772 | 56,442 | 160,215 | 2,571,654 |
| chr4 | 19,102,811 | 6,016,599 | 1,460,219 | 1,973,905 | 647,314 | 36725 | 980,528 | 4,837,136 | 55,124 | 161,166 | 2,688,914 |
| chr5 | 18,819,841 | 5,600,366 | 1,498,886 | 1,999,842 | 574,644 | 39951 | 1,280,709 | 5,183,157 | 55,073 | 154,616 | 2,201,669 |
| chr6 | 18,788,989 | 7,125,284 | 1,145,162 | 1,341,896 | 713,567 | 42130 | 1,069,703 | 4,233,351 | 63,831 | 171,265 | 2,626,420 |
| chr7 | 18,392,816 | 5,871,751 | 1,272,382 | 1,673,093 | 563,108 | 57647 | 1,639,155 | 5,174,895 | 56,877 | 154,456 | 1,730,173 |
| chr8 | 18,352,810 | 6,708,116 | 1,021,656 | 1,326,943 | 729,659 | 43451 | 1,280,179 | 4,672,372 | 58,199 | 156,033 | 2,121,847 |
| chr9 | 18,070,974 | 5,383,751 | 1,333,369 | 1,917,235 | 580,773 | 83571 | 945,043 | 5,151,506 | 51,007 | 150,173 | 2,309,239 |
| chr10 | 17,741,200 | 6,644,184 | 948,652 | 1,579,690 | 476,322 | 60966 | 988,495 | 3,919,245 | 57,012 | 148,721 | 2,713,397 |
| chr11 | 17,282,498 | 5,125,834 | 1,241,894 | 2,030,548 | 480,201 | 39350 | 1,134,080 | 4,858,542 | 49,115 | 139,685 | 1,992,233 |
| chr12 | 16,740,881 | 4,704,818 | 1,062,585 | 1,463,011 | 660,817 | 41766 | 2,333,080 | 5,480,209 | 47,168 | 128,047 | 642,518 |
| chr13 | 15,436,925 | 6,284,970 | 720,849 | 1,280,104 | 441,757 | 55115 | 790,315 | 3,167,138 | 48,862 | 130,717 | 2,330,875 |
| chr14 | 15,407,298 | 4,378,734 | 964,426 | 1,699,245 | 462,606 | 38892 | 508,801 | 4,358,665 | 47,791 | 125,944 | 2,649,952 |
| chr15 | 14,069,062 | 4,370,540 | 817,124 | 1,308,589 | 460,946 | 66555 | 1,745,424 | 4,062,564 | 43,679 | 111,702 | 949,907 |
| chr16 | 12,746,868 | 4,215,633 | 781,546 | 1,121,035 | 335,132 | 77555 | 783,765 | 3,503,990 | 38,460 | 104,412 | 1,659,580 |
| Total | 288,380,347 | 96,322,343 | 18,546,981 | 26,269,609 | 9,499,657 | 838,253 | 20,051,267 | 74,804,697 | 870,062 | 2,388,885 | 38,870,394 |
| Percentage (%) | 100 | 33.40 | 6.43 | 9.11 | 3.29 | 0.29 | 6.95 | 25.94 | 0.30 | 0.83 | 13.48 |

**Table S3.** Functional annotation of the 448 *Spondias tuberosa* genes in the 97 gene families significantly expanded in *S. tuberosa* (CAFE5, $P < 0.05$; OrthoVenn3, seven species).

| Gene ID | Cluster | Function | GO IDs | GO Names |
|---|---|---|---|---|
| tube_chr3_001128.1 | cluster22 | disease resistance protein At4g27190-like isoform X1 | P:GO:0006952; P:GO:0098542; F:GO:0016887; F:GO:0043531 | P:defense response; P:defense response to other organism; F:ATP hydrolysis activity; F:ADP binding |
| tube_chr16_001134.1 | cluster22 | probable disease resistance protein At4g27220 isoform X1 | P:GO:0006952; P:GO:0098542; F:GO:0016887; F:GO:0043531 | P:defense response; P:defense response to other organism; F:ATP hydrolysis activity; F:ADP binding |
| tube_chr16_000934.1 | cluster22 | probable disease resistance protein At4g27220 | P:GO:0006952; P:GO:0098542; F:GO:0016887; F:GO:0043531 | P:defense response; P:defense response to other organism; F:ATP hydrolysis activity; F:ADP binding |
| tube_chr16_000930.1 | cluster22 | disease resistance protein At4g27190-like isoform X1 | P:GO:0006952; P:GO:0098542; F:GO:0016887; F:GO:0043531 | P:defense response; P:defense response to other organism; F:ATP hydrolysis activity; F:ADP binding |
| tube_chr16_000927.1 | cluster22 | disease resistance protein At4g27190-like isoform X1 | P:GO:0006952; P:GO:0098542; F:GO:0016887; F:GO:0043531 | P:defense response; P:defense response to other organism; F:ATP hydrolysis activity; F:ADP binding |
| tube_chr15_000983.1 | cluster22 | disease resistance protein At4g27190-like isoform X1 | P:GO:0006952; P:GO:0098542; F:GO:0016887; F:GO:0043531 | P:defense response; P:defense response to other organism; F:ATP hydrolysis activity; F:ADP binding |
| tube_chr15_000981.1 | cluster22 | disease resistance protein At4g27190-like isoform X1 | P:GO:0006952; P:GO:0098542; F:GO:0016887; F:GO:0043531 | P:defense response; P:defense response to other organism; F:ATP hydrolysis activity; F:ADP binding |
| tube_chr15_000980.1 | cluster22 | disease resistance protein At4g27190-like isoform X1 | P:GO:0006952; P:GO:0098542; F:GO:0016887; F:GO:0043531 | P:defense response; P:defense response to other organism; F:ATP hydrolysis activity; F:ADP binding |
| tube_chr15_000978.1 | cluster22 | disease resistance protein At4g27190-like isoform X1 | P:GO:0006952; P:GO:0098542; F:GO:0016887; F:GO:0043531 | P:defense response; P:defense response to other organism; F:ATP hydrolysis activity; F:ADP binding |
| tube_chr15_000977.1 | cluster22 | disease resistance protein At4g27190-like isoform X1 | P:GO:0006952; P:GO:0098542; F:GO:0016887; F:GO:0043531 | P:defense response; P:defense response to other organism; F:ATP hydrolysis activity; F:ADP binding |
| tube_chr16_001130.1 | cluster22 | disease resistance protein At4g27190-like isoform X1 | | |
| tube_chr16_000063.1 | cluster22 | disease resistance protein At4g27190-like isoform X1 | P:GO:0006952; P:GO:0098542; F:GO:0016887; F:GO:0043531 | P:defense response; P:defense response to other organism; F:ATP hydrolysis activity; F:ADP binding |
| tube_chr16_000064.1 | cluster22 | disease resistance protein At4g27190-like isoform X1 | P:GO:0006952; P:GO:0098542; F:GO:0016887; F:GO:0043531 | P:defense response; P:defense response to other organism; F:ATP hydrolysis activity; F:ADP binding |
| tube_chr16_000925.1 | cluster22 | disease resistance protein At4g27190-like isoform X1 | P:GO:0006952; P:GO:0098542; F:GO:0016887; F:GO:0043531 | P:defense response; P:defense response to other organism; F:ATP hydrolysis activity; F:ADP binding |
| tube_chr16_001131.1 | cluster22 | disease resistance protein At4g27190-like isoform X1 | P:GO:0006952; P:GO:0098542; F:GO:0016887; F:GO:0043531 | P:defense response; P:defense response to other organism; F:ATP hydrolysis activity; F:ADP binding |
| tube_chr14_000749.1 | cluster48 | probable disease resistance protein At5g63020 | P:GO:0006952 | P:defense response |
| tube_chr14_000489.1 | cluster48 | probable disease resistance protein At5g63020 | P:GO:0006952 | P:defense response |
| tube_chr14_000354.1 | cluster48 | probable disease resistance protein At5g63020 isoform X1 | P:GO:0006952 | P:defense response |
| tube_chr14_000350.1 | cluster48 | probable disease resistance protein At5g63020 | P:GO:0006952 | P:defense response |
| tube_chr14_000347.1 | cluster48 | probable disease resistance protein At5g63020 | P:GO:0006952 | P:defense response |
| tube_chr14_000345.1 | cluster48 | probable disease resistance protein At5g63020 | P:GO:0006952 | P:defense response |
| tube_chr14_000344.1 | cluster48 | probable disease resistance protein At5g63020 | P:GO:0006952 | P:defense response |
| tube_chr14_000293.1 | cluster48 | probable disease resistance protein At5g63020 | P:GO:0006952 | P:defense response |
| tube_chr14_000292.1 | cluster48 | probable disease resistance protein At5g63020 | P:GO:0006952 | P:defense response |
| tube_chr14_000349.1 | cluster48 | probable disease resistance protein At5g63020 | P:GO:0006952 | P:defense response |
| tube_chr14_000352.1 | cluster48 | probable disease resistance protein At5g63020 | P:GO:0006952 | P:defense response |
| tube_chr9_000666.1 | cluster79 | subtilisin-like protease SBT5.4 | P:GO:0006508; P:GO:0009610; F:GO:0004252; C:GO:0005615 | P:proteolysis; P:response to symbiotic fungus; F:serine-type endopeptidase activity; C:obsolete extracellular space |
| tube_chr1_001657.1 | cluster79 | subtilisin-like protease SBT5.3 | P:GO:0006508; F:GO:0004252 | P:proteolysis; F:serine-type endopeptidase activity |
| tube_chr1_001535.1 | cluster79 | subtilisin-like protease SBT5.3 | P:GO:0006508; F:GO:0004252; C:GO:0005615 | P:proteolysis; F:serine-type endopeptidase activity; C:obsolete extracellular space |
| tube_chr7_000947.1 | cluster79 | subtilisin-like protease SBT5.4 | F:GO:0004252; C:GO:0005615 | F:serine-type endopeptidase activity; C:obsolete extracellular space |
| tube_chr5_001337.1 | cluster79 | subtilisin-like protease SBT5.4 | F:GO:0004252; C:GO:0005615 | F:serine-type endopeptidase activity; C:obsolete extracellular space |
| tube_chr5_001336.1 | cluster79 | subtilisin-like protease SBT5.4 | F:GO:0004252; C:GO:0005615 | F:serine-type endopeptidase activity; C:obsolete extracellular space |
| tube_chr5_000096.1 | cluster79 | subtilisin-like protease SBT5.4 | F:GO:0004252; C:GO:0005615 | F:serine-type endopeptidase activity; C:obsolete extracellular space |

| Gene ID | Cluster | Function | GO IDs | GO Names |
|---|---|---|---|---|
| tube_chr16_000596.1 | cluster79 | subtilisin-like protease SBT5.4 | F:GO:0004252; C:GO:0005615 | F:serine-type endopeptidase activity; C:obsolete extracellular space |
| tube_chr16_000595.1 | cluster79 | subtilisin-like protease SBT5.4 | F:GO:0004252; C:GO:0005615 | F:serine-type endopeptidase activity; C:obsolete extracellular space |
| tube_chr16_000594.1 | cluster79 | subtilisin-like protease SBT5.4 | F:GO:0004252; C:GO:0005615 | F:serine-type endopeptidase activity; C:obsolete extracellular space |
| tube_chr16_000593.1 | cluster79 | subtilisin-like protease SBT5.4 | F:GO:0004252; C:GO:0005615 | F:serine-type endopeptidase activity; C:obsolete extracellular space |
| tube_chr16_000592.1 | cluster79 | subtilisin-like protease SBT5.4 | F:GO:0004252; C:GO:0005615 | F:serine-type endopeptidase activity; C:obsolete extracellular space |
| tube_chr16_000590.1 | cluster79 | subtilisin-like protease SBT5.4 | F:GO:0008236; C:GO:0005576 | F:serine-type peptidase activity; C:extracellular region |
| tube_chr15_000903.1 | cluster79 | subtilisin-like protease SBT5.4 | F:GO:0004252; C:GO:0005615 | F:serine-type endopeptidase activity; C:obsolete extracellular space |
| tube_chr5_001339.1 | cluster79 | subtilisin-like protease SBT5.4 | F:GO:0008236 | F:serine-type peptidase activity |
| tube_chr16_000591.1 | cluster79 | subtilisin-like protease SBT5.4 | F:GO:0008236; C:GO:0005576 | F:serine-type peptidase activity; C:extracellular region |
| tube_chr11_000486.1 | cluster83 | germin-like protein subfamily 1 member 13 | | |
| tube_chr11_000298.1 | cluster83 | germin-like protein subfamily 1 member 17 | | |
| tube_chr11_000299.1 | cluster83 | germin-like protein subfamily 1 member 17 | | |
| tube_chr11_000488.1 | cluster83 | germin-like protein subfamily 1 member 17 | | |
| tube_chr11_000490.1 | cluster83 | germin-like protein subfamily 1 member 17 | | |
| tube_chr11_000877.1 | cluster83 | germin-like protein subfamily 1 member 13 | | |
| tube_chr11_000884.1 | cluster83 | germin-like protein subfamily 1 member 13 | | |
| tube_chr11_000886.1 | cluster83 | germin-like protein subfamily 1 member 13 | | |
| tube_chr11_000931.1 | cluster83 | germin-like protein subfamily 1 member 17 | | |
| tube_chr11_000885.1 | cluster83 | germin-like protein subfamily 1 member 13 | F:GO:0030145; C:GO:0048046 | F:manganese ion binding; C:apoplast |
| tube_chr11_000735.1 | cluster83 | germin-like protein subfamily 1 member 17 | | |
| tube_chr11_000736.1 | cluster83 | germin-like protein subfamily 1 member 17 | | |
| tube_chr6_000815.1 | cluster106 | cysteine-rich receptor-like protein kinase 19 | P:GO:0005975; P:GO:0006955; P:GO:0007165; F:GO:0004674; F:GO:0005524; F:GO:0008061; C:GO:0005886 | P:carbohydrate metabolic process; P:immune response; P:signal transduction; F:protein serine/threonine kinase activity; F:ATP binding; F:chitin binding; C:plasma membrane |
| tube_chr6_000811.1 | cluster106 | receptor-like serine/threonine-protein kinase SD1-8 | P:GO:0005975; F:GO:0004674; F:GO:0005524; F:GO:0008061; F:GO:0009055; C:GO:0005886 | P:carbohydrate metabolic process; F:protein serine/threonine kinase activity; F:ATP binding; F:chitin binding; F:electron transfer activity; C:plasma membrane |
| tube_chr6_000808.1 | cluster106 | receptor-like serine/threonine-protein kinase SD1-8 | | |
| tube_chr6_000798.1 | cluster106 | receptor-like serine/threonine-protein kinase SD1-8 | P:GO:0005975; P:GO:0006032; F:GO:0004568; F:GO:0004672; F:GO:0008061; C:GO:0005576; C:GO:0016020 | P:carbohydrate metabolic process; P:chitin catabolic process; F:chitinase activity; F:protein kinase activity; F:chitin binding; C:extracellular region; C:membrane |
| tube_chr6_000737.1 | cluster106 | Receptor-like serine/threonine-protein kinase | P:GO:0005975; P:GO:0006955; P:GO:0007165; F:GO:0004674; F:GO:0005524; F:GO:0008061; C:GO:0005886 | P:carbohydrate metabolic process; P:immune response; P:signal transduction; F:protein serine/threonine kinase activity; F:ATP binding; F:chitin binding; C:plasma membrane |
| tube_chr6_000735.1 | cluster106 | Receptor-like serine/threonine-protein kinase | P:GO:0005975; P:GO:0006955; P:GO:0007165; F:GO:0004674; F:GO:0005524; F:GO:0008061; C:GO:0005886 | P:carbohydrate metabolic process; P:immune response; P:signal transduction; F:protein serine/threonine kinase activity; F:ATP binding; F:chitin binding; C:plasma membrane |
| tube_chr6_000734.1 | cluster106 | Receptor-like serine/threonine-protein kinase | P:GO:0005975; P:GO:0006955; P:GO:0007165; F:GO:0004674; F:GO:0005524; F:GO:0008061; C:GO:0005886 | P:carbohydrate metabolic process; P:immune response; P:signal transduction; F:protein serine/threonine kinase activity; F:ATP binding; F:chitin binding; C:plasma membrane |
| tube_chr6_000733.1 | cluster106 | cysteine-rich receptor protein kinase | P:GO:0009987; P:GO:0050896; F:GO:0004672; F:GO:0097367; C:GO:0016020 | P:cellular process; P:response to stimulus; F:protein kinase activity; F:carbohydrate derivative binding; C:membrane |
| tube_chr8_000077.1 | cluster125 | MDIS1-interacting receptor like kinase 2-like | P:GO:0009755; F:GO:0004672; F:GO:0004674; F:GO:0005524; F:GO:0038023; C:GO:0005886; C:GO:0016020 | P:hormone-mediated signaling pathway; F:protein kinase activity; F:protein serine/threonine kinase activity; F:ATP binding; F:signaling receptor activity; C:plasma membrane; C:membrane |
| tube_chr8_001484.1 | cluster125 | MDIS1-interacting receptor like kinase 2 | F:GO:0004674; F:GO:0005524; C:GO:0005886 | F:protein serine/threonine kinase activity; F:ATP binding; C:plasma membrane |
| tube_chr8_000074.1 | cluster125 | protein kinase domain-containing protein | P:GO:0009755; F:GO:0038023; C:GO:0005886 | P:hormone-mediated signaling pathway; F:signaling receptor activity; C:plasma membrane |
| tube_chr8_000076.1 | cluster125 | putative leucine-rich repeat receptor-like protein kinase | F:GO:0005488 | F:binding |
| tube_chr8_001485.1 | cluster125 | MDIS1-interacting receptor like kinase 2 | F:GO:0004672; F:GO:0004674; F:GO:0005524; C:GO:0016020 | F:protein kinase activity; F:protein serine/threonine kinase activity; F:ATP binding; C:membrane |
| tube_chr8_001486.1 | cluster125 | MDIS1-interacting receptor like kinase 2 | F:GO:0004672; F:GO:0004674; F:GO:0005524; C:GO:0005886; C:GO:0016020 | F:protein kinase activity; F:protein serine/threonine kinase activity; F:ATP binding; C:plasma membrane; C:membrane |

| Gene ID | Cluster | Function | GO IDs | GO Names |
|---|---|---|---|---|
| tube_chr1_001437.1 | cluster126 | cellulose synthase-like protein B4 | P:GO:0030244; P:GO:0071555; F:GO:0016760; C:GO:0012505; C:GO:0016020 | P:cellulose biosynthetic process; P:cell wall organization; F:cellulose synthase (UDP-forming) activity; C:endomembrane system; C:membrane |
| tube_chr1_001436.1 | cluster126 | cellulose synthase-like protein B4 | P:GO:0030244; P:GO:0071555; F:GO:0016760; C:GO:0012505; C:GO:0016020 | P:cellulose biosynthetic process; P:cell wall organization; F:cellulose synthase (UDP-forming) activity; C:endomembrane system; C:membrane |
| tube_chr1_001435.1 | cluster126 | cellulose synthase-like protein B4 | P:GO:0030244; P:GO:0071555; F:GO:0016760; C:GO:0012505; C:GO:0016020 | P:cellulose biosynthetic process; P:cell wall organization; F:cellulose synthase (UDP-forming) activity; C:endomembrane system; C:membrane |
| tube_chr1_001434.1 | cluster126 | cellulose synthase-like protein B4 | P:GO:0030244; P:GO:0071555; F:GO:0016760; C:GO:0012505; C:GO:0016020 | P:cellulose biosynthetic process; P:cell wall organization; F:cellulose synthase (UDP-forming) activity; C:endomembrane system; C:membrane |
| tube_chr1_001433.1 | cluster126 | cellulose synthase-like protein B4 | P:GO:0030244; P:GO:0071555; F:GO:0016760; C:GO:0012505; C:GO:0016020 | P:cellulose biosynthetic process; P:cell wall organization; F:cellulose synthase (UDP-forming) activity; C:endomembrane system; C:membrane |
| tube_chr1_001430.1 | cluster126 | cellulose synthase-like protein B4 | P:GO:0030244; P:GO:0071555; F:GO:0016760; C:GO:0012505; C:GO:0016020 | P:cellulose biosynthetic process; P:cell wall organization; F:cellulose synthase (UDP-forming) activity; C:endomembrane system; C:membrane |
| tube_chr1_001426.1 | cluster126 | cellulose synthase-like protein B4 | P:GO:0030244; P:GO:0071555; F:GO:0016760; C:GO:0012505; C:GO:0016020 | P:cellulose biosynthetic process; P:cell wall organization; F:cellulose synthase (UDP-forming) activity; C:endomembrane system; C:membrane |
| tube_chr1_001422.1 | cluster126 | cellulose synthase-like protein B4 | P:GO:0030244; P:GO:0071555; F:GO:0016760; C:GO:0012505; C:GO:0016020 | P:cellulose biosynthetic process; P:cell wall organization; F:cellulose synthase (UDP-forming) activity; C:endomembrane system; C:membrane |
| tube_chr1_001421.1 | cluster126 | cellulose synthase-like protein B4 | P:GO:0030244; P:GO:0071555; F:GO:0016760; C:GO:0012505; C:GO:0016020 | P:cellulose biosynthetic process; P:cell wall organization; F:cellulose synthase (UDP-forming) activity; C:endomembrane system; C:membrane |
| tube_chr1_001420.1 | cluster126 | cellulose synthase-like protein B4 | P:GO:0030244; P:GO:0071555; F:GO:0016760; C:GO:0012505; C:GO:0016020 | P:cellulose biosynthetic process; P:cell wall organization; F:cellulose synthase (UDP-forming) activity; C:endomembrane system; C:membrane |
| tube_chr9_000779.1 | cluster129 | probable F-box protein At4g22030 | | |
| tube_chr9_000778.1 | cluster129 | probable F-box protein At4g22030 | | |
| tube_chr9_000780.1 | cluster129 | probable F-box protein At4g22030 | | |
| tube_chr9_000781.1 | cluster129 | probable F-box protein At4g22030 | | |
| tube_chr9_000782.1 | cluster129 | probable F-box protein At4g22030 | | |
| tube_chr9_000783.1 | cluster129 | probable F-box protein At4g22030 | | |
| tube_chr14_000182.1 | cluster142 | putative leucine-rich repeat receptor-like serine/threonine-protein kinase At2g19230 | F:GO:0004672 | F:protein kinase activity |
| tube_chr14_000181.1 | cluster142 | putative leucine-rich repeat receptor-like serine/threonine-protein kinase At2g19230 | F:GO:0004672 | F:protein kinase activity |
| tube_chr14_000180.1 | cluster142 | putative leucine-rich repeat receptor-like serine/threonine-protein kinase At2g19230 | F:GO:0004672 | F:protein kinase activity |
| tube_chr14_000179.1 | cluster142 | putative leucine-rich repeat receptor-like serine/threonine-protein kinase At2g19230 | F:GO:0004672 | F:protein kinase activity |
| tube_chr14_000177.1 | cluster142 | putative leucine-rich repeat receptor-like serine/threonine-protein kinase At2g19230 | F:GO:0004672 | F:protein kinase activity |
| tube_chr14_000527.1 | cluster142 | putative leucine-rich repeat receptor-like serine/threonine-protein kinase At2g19230 | F:GO:0004672 | F:protein kinase activity |
| tube_chr5_000502.1 | cluster151 | putative polyol transporter 1 | P:GO:0034219; P:GO:1902600; F:GO:0005351; C:GO:0016020 | P:carbohydrate transmembrane transport; P:proton transmembrane transport; F:carbohydrate:proton symporter activity; C:membrane |
| tube_chr5_000503.1 | cluster151 | putative polyol transporter 1 | P:GO:0034219; P:GO:1902600; F:GO:0005351; C:GO:0016020 | P:carbohydrate transmembrane transport; P:proton transmembrane transport; F:carbohydrate:proton symporter activity; C:membrane |
| tube_chr5_000504.1 | cluster151 | putative polyol transporter 1 | P:GO:0034219; P:GO:1902600; F:GO:0005351; C:GO:0016020 | P:carbohydrate transmembrane transport; P:proton transmembrane transport; F:carbohydrate:proton symporter activity; C:membrane |
| tube_chr5_000505.1 | cluster151 | putative polyol transporter 1 | P:GO:0034219; P:GO:1902600; F:GO:0005351; C:GO:0016020 | P:carbohydrate transmembrane transport; P:proton transmembrane transport; F:carbohydrate:proton symporter activity; C:membrane |
| tube_chr10_001420.1 | cluster190 | gibberellin 20 oxidase 2-like | F:GO:0016491 | F:oxidoreductase activity |
| tube_chr10_000139.1 | cluster190 | gibberellin 20 oxidase 2-like | P:GO:0009416; P:GO:0009686; P:GO:0009826; P:GO:0009908; F:GO:0045544 | P:response to light stimulus; P:gibberellin biosynthetic process; P:unidimensional cell growth; P:flower development; F:gibberellin 20-oxidase activity |
| tube_chr10_001654.1 | cluster190 | gibberellin 20 oxidase 2-like | P:GO:0009416; P:GO:0009686; P:GO:0009826; P:GO:0009908; F:GO:0045544 | P:response to light stimulus; P:gibberellin biosynthetic process; P:unidimensional cell growth; P:flower development; F:gibberellin 20-oxidase activity |
| tube_chr10_001656.1 | cluster190 | gibberellin 20 oxidase 2-like | P:GO:0009416; P:GO:0009686; P:GO:0009826; P:GO:0009908; F:GO:0045544 | P:response to light stimulus; P:gibberellin biosynthetic process; P:unidimensional cell growth; P:flower development; F:gibberellin 20-oxidase activity |

| Gene ID | Cluster | Function | GO IDs | GO Names |
|---|---|---|---|---|
| tube_chr14_000468.1 | cluster190 | gibberellin 20 oxidase 2-like | P:GO:0009416; P:GO:0009686; P:GO:0009826; P:GO:0009908; F:GO:0045544 | P:response to light stimulus; P:gibberellin biosynthetic process; P:unidimensional cell growth; P:flower development; F:gibberellin 20-oxidase activity |
| tube_chr14_000469.1 | cluster190 | gibberellin 20 oxidase 2-like | P:GO:0009416; P:GO:0009686; P:GO:0009826; P:GO:0009908; F:GO:0045544 | P:response to light stimulus; P:gibberellin biosynthetic process; P:unidimensional cell growth; P:flower development; F:gibberellin 20-oxidase activity |
| tube_chr14_000470.1 | cluster190 | gibberellin 20 oxidase 1-B-like | P:GO:0009987; P:GO:0048856; F:GO:0016705; F:GO:0051213 | P:cellular process; P:anatomical structure development; F:oxidoreductase activity, acting on paired donors, with incorporation or reduction of molecular oxygen; F:dioxygenase activity |
| tube_chr14_000471.1 | cluster190 | gibberellin 20 oxidase 2-like | P:GO:0009416; P:GO:0009686; P:GO:0009826; P:GO:0009908; F:GO:0045544 | P:response to light stimulus; P:gibberellin biosynthetic process; P:unidimensional cell growth; P:flower development; F:gibberellin 20-oxidase activity |
| tube_chr14_000472.1 | cluster190 | gibberellin 20 oxidase 2-like | P:GO:0009416; P:GO:0009686; P:GO:0009826; P:GO:0009908; F:GO:0045544 | P:response to light stimulus; P:gibberellin biosynthetic process; P:unidimensional cell growth; P:flower development; F:gibberellin 20-oxidase activity |
| tube_chr14_000473.1 | cluster190 | gibberellin 20 oxidase 2-like | P:GO:0009416; P:GO:0009686; P:GO:0009826; P:GO:0009908; F:GO:0045544 | P:response to light stimulus; P:gibberellin biosynthetic process; P:unidimensional cell growth; P:flower development; F:gibberellin 20-oxidase activity |
| tube_chr14_000474.1 | cluster190 | gibberellin 20 oxidase 2-like | P:GO:0009416; P:GO:0009686; P:GO:0009826; P:GO:0009908; F:GO:0045544 | P:response to light stimulus; P:gibberellin biosynthetic process; P:unidimensional cell growth; P:flower development; F:gibberellin 20-oxidase activity |
| tube_chr14_000656.1 | cluster190 | gibberellin 20 oxidase 2-like | P:GO:0009416; P:GO:0009686; P:GO:0009826; P:GO:0009908; F:GO:0045544 | P:response to light stimulus; P:gibberellin biosynthetic process; P:unidimensional cell growth; P:flower development; F:gibberellin 20-oxidase activity |
| tube_chr14_000657.1 | cluster190 | gibberellin 20 oxidase 2-like | P:GO:0009416; P:GO:0009686; P:GO:0009826; P:GO:0009908; F:GO:0045544 | P:response to light stimulus; P:gibberellin biosynthetic process; P:unidimensional cell growth; P:flower development; F:gibberellin 20-oxidase activity |
| tube_chr14_000669.1 | cluster190 | gibberellin 20 oxidase 2-like | F:GO:0016491 | F:oxidoreductase activity |
| tube_chr4_000040.1 | cluster190 | gibberellin 20 oxidase 2-like | P:GO:0009416; P:GO:0009686; P:GO:0009826; P:GO:0009908; F:GO:0045544 | P:response to light stimulus; P:gibberellin biosynthetic process; P:unidimensional cell growth; P:flower development; F:gibberellin 20-oxidase activity |
| tube_chr7_000218.1 | cluster190 | gibberellin 20 oxidase 2-like | P:GO:0009416; P:GO:0009686; P:GO:0009826; P:GO:0009908; F:GO:0045544 | P:response to light stimulus; P:gibberellin biosynthetic process; P:unidimensional cell growth; P:flower development; F:gibberellin 20-oxidase activity |
| tube_chr7_000219.1 | cluster190 | gibberellin 20 oxidase 2-like | P:GO:0009416; P:GO:0009686; P:GO:0009826; P:GO:0009908; F:GO:0045544 | P:response to light stimulus; P:gibberellin biosynthetic process; P:unidimensional cell growth; P:flower development; F:gibberellin 20-oxidase activity |
| tube_chr7_000220.1 | cluster190 | gibberellin 20 oxidase 2-like | P:GO:0009416; P:GO:0009686; P:GO:0009826; P:GO:0009908; F:GO:0045544 | P:response to light stimulus; P:gibberellin biosynthetic process; P:unidimensional cell growth; P:flower development; F:gibberellin 20-oxidase activity |
| tube_chr7_000221.1 | cluster190 | gibberellin 20 oxidase 2-like | P:GO:0009416; P:GO:0009686; P:GO:0009826; P:GO:0009908; F:GO:0045544 | P:response to light stimulus; P:gibberellin biosynthetic process; P:unidimensional cell growth; P:flower development; F:gibberellin 20-oxidase activity |
| tube_chr7_000222.1 | cluster190 | gibberellin 20 oxidase 2-like | P:GO:0009416; P:GO:0009686; P:GO:0009826; P:GO:0009908; F:GO:0045544 | P:response to light stimulus; P:gibberellin biosynthetic process; P:unidimensional cell growth; P:flower development; F:gibberellin 20-oxidase activity |
| tube_chr7_000223.1 | cluster190 | gibberellin 20 oxidase 2-like | P:GO:0009416; P:GO:0009686; P:GO:0009826; P:GO:0009908; F:GO:0045544 | P:response to light stimulus; P:gibberellin biosynthetic process; P:unidimensional cell growth; P:flower development; F:gibberellin 20-oxidase activity |
| tube_chr9_000598.1 | cluster197 | LRR receptor-like kinase | F:GO:0004672; F:GO:0004674; F:GO:0005524; C:GO:0016020 | F:protein kinase activity; F:protein serine/threonine kinase activity; F:ATP binding; C:membrane |
| tube_chr9_000594.1 | cluster197 | LRR receptor-like kinase | F:GO:0004672 | F:protein kinase activity |
| tube_chr9_000593.1 | cluster197 | LRR receptor-like kinase | F:GO:0004672; F:GO:0004674; F:GO:0005524; C:GO:0016020 | F:protein kinase activity; F:protein serine/threonine kinase activity; F:ATP binding; C:membrane |
| tube_chr9_000595.1 | cluster197 | LRR receptor-like kinase | F:GO:0004672; F:GO:0004674; F:GO:0005524; C:GO:0016020 | F:protein kinase activity; F:protein serine/threonine kinase activity; F:ATP binding; C:membrane |
| tube_chr9_000599.1 | cluster197 | LRR receptor-like kinase | F:GO:0004672; F:GO:0004674; F:GO:0005524; C:GO:0016020 | F:protein kinase activity; F:protein serine/threonine kinase activity; F:ATP binding; C:membrane |
| tube_chr9_000600.1 | cluster197 | LRR receptor-like kinase | F:GO:0004672; F:GO:0004674; F:GO:0005524; C:GO:0016020 | F:protein kinase activity; F:protein serine/threonine kinase activity; F:ATP binding; C:membrane |
| tube_chr9_000602.1 | cluster197 | LRR receptor-like kinase | F:GO:0004672; F:GO:0004674; F:GO:0005524; C:GO:0016020 | F:protein kinase activity; F:protein serine/threonine kinase activity; F:ATP binding; C:membrane |
| tube_chr9_000603.1 | cluster197 | LRR receptor-like kinase | F:GO:0004672; F:GO:0004674; F:GO:0005524; C:GO:0016020 | F:protein kinase activity; F:protein serine/threonine kinase activity; F:ATP binding; C:membrane |
| tube_chr9_000596.1 | cluster197 | putative LRR receptor-like serine/threonine-protein kinase | | |
| tube_chr9_000285.1 | cluster209 | d-limonene synthase | P:GO:0016102; F:GO:0000287; F:GO:0010333 | P:diterpenoid biosynthetic process; F:magnesium ion binding; F:terpene synthase activity |
| tube_chr9_001185.1 | cluster209 | (R)-limonene synthase 1, chloroplastic-like | P:GO:0016102; F:GO:0000287; F:GO:0010333 | P:diterpenoid biosynthetic process; F:magnesium ion binding; F:terpene synthase activity |
| tube_chr9_001137.1 | cluster209 | (R)-limonene synthase 1, chloroplastic-like | P:GO:0016102; F:GO:0000287; F:GO:0010333 | P:diterpenoid biosynthetic process; F:magnesium ion binding; F:terpene synthase activity |

| Gene ID | Cluster | Function | GO IDs | GO Names |
|---|---|---|---|---|
| tube_chr9_001186.1 | cluster209 | (R)-limonene synthase 1, chloroplastic-like | P:GO:0016102; F:GO:0000287; F:GO:0010333 | P:diterpenoid biosynthetic process; F:magnesium ion binding; F:terpene synthase activity |
| tube_chr9_001187.1 | cluster209 | (R)-limonene synthase 1, chloroplastic-like | P:GO:0016102; F:GO:0000287; F:GO:0010333 | P:diterpenoid biosynthetic process; F:magnesium ion binding; F:terpene synthase activity |
| tube_chr16_000853.1 | cluster217 | (-)-germacrene D synthase-like | P:GO:0016102; F:GO:0000287; F:GO:0010333 | P:diterpenoid biosynthetic process; F:magnesium ion binding; F:terpene synthase activity |
| tube_chr16_000856.1 | cluster217 | alpha-humulene/(-)-(E)-beta-caryophyllene synthase | P:GO:0016102; F:GO:0000287; F:GO:0010333 | P:diterpenoid biosynthetic process; F:magnesium ion binding; F:terpene synthase activity |
| tube_chr16_000860.1 | cluster217 | alpha-humulene/(-)-(E)-beta-caryophyllene synthase | P:GO:0016102; F:GO:0000287; F:GO:0010333 | P:diterpenoid biosynthetic process; F:magnesium ion binding; F:terpene synthase activity |
| tube_chr16_000554.1 | cluster217 | alpha-humulene/(-)-(E)-beta-caryophyllene synthase | P:GO:0016102; F:GO:0000287; F:GO:0010333 | P:diterpenoid biosynthetic process; F:magnesium ion binding; F:terpene synthase activity |
| tube_chr16_000168.1 | cluster217 | alpha-humulene/(-)-(E)-beta-caryophyllene synthase | P:GO:0016102; F:GO:0000287; F:GO:0010333 | P:diterpenoid biosynthetic process; F:magnesium ion binding; F:terpene synthase activity |
| tube_chr16_000167.1 | cluster217 | alpha-humulene/(-)-(E)-beta-caryophyllene synthase | P:GO:0016102; F:GO:0000287; F:GO:0010333 | P:diterpenoid biosynthetic process; F:magnesium ion binding; F:terpene synthase activity |
| tube_chr16_000166.1 | cluster217 | alpha-humulene/(-)-(E)-beta-caryophyllene synthase | P:GO:0016102; F:GO:0000287; F:GO:0010333 | P:diterpenoid biosynthetic process; F:magnesium ion binding; F:terpene synthase activity |
| tube_chr5_000763.1 | cluster229 | uncharacterized protein LOC123213678 | P:GO:0048731 | P:system development |
| tube_chr5_000762.1 | cluster229 | uncharacterized protein LOC123227776 | P:GO:0048731 | P:system development |
| tube_chr5_000760.1 | cluster229 | uncharacterized protein LOC116106739 | P:GO:0048364; P:GO:0048367 | P:root development; P:shoot system development |
| tube_chr5_000761.1 | cluster229 | uncharacterized protein LOC123213678 | P:GO:0048364; P:GO:0048367 | P:root development; P:shoot system development |
| tube_chr7_000602.1 | cluster233 | senescence-associated protein 13-like | F:GO:0016491 | F:oxidoreductase activity |
| tube_chr7_000605.1 | cluster233 | senescence-associated protein 13-like | F:GO:0016491 | F:oxidoreductase activity |
| tube_chr7_000606.1 | cluster233 | tropinone reductase homolog At2g29290-like | F:GO:0016491 | F:oxidoreductase activity |
| tube_chr7_000603.1 | cluster233 | senescence-associated protein 13-like | F:GO:0016491 | F:oxidoreductase activity |
| tube_chr7_000599.1 | cluster233 | F-box domain-containing protein | F:GO:0016491 | F:oxidoreductase activity |
| tube_chr7_000608.1 | cluster233 | SDR family oxidoreductase | F:GO:0016491 | F:oxidoreductase activity |
| tube_chr15_000398.1 | cluster241 | PROPROTEIN CONVERTASE SUBTILISIN/KEXIN | P:GO:0006508; F:GO:0004252; C:GO:0005576; C:GO:0016020 | P:proteolysis; F:serine-type endopeptidase activity; C:extracellular region; C:membrane |
| tube_chr15_000397.1 | cluster241 | subtilisin-like protease SBT4.3 | P:GO:0006508; F:GO:0004252; C:GO:0016020 | P:proteolysis; F:serine-type endopeptidase activity; C:membrane |
| tube_chr15_000396.1 | cluster241 | cucumisin-like | P:GO:0006508; F:GO:0004252 | P:proteolysis; F:serine-type endopeptidase activity |
| tube_chr15_000395.1 | cluster241 | cucumisin-like | P:GO:0006508; F:GO:0004252 | P:proteolysis; F:serine-type endopeptidase activity |
| tube_chr12_001215.1 | cluster266 | anthocyanidin 3-O-glucosyltransferase 2-like | | |
| tube_chr12_001211.1 | cluster266 | anthocyanidin 3-O-glucosyltransferase 2-like | | |
| tube_chr12_001212.1 | cluster266 | anthocyanidin 3-O-glucosyltransferase 2-like | | |
| tube_chr12_001213.1 | cluster266 | anthocyanidin 3-O-glucosyltransferase 2-like | | |
| tube_chr12_001214.1 | cluster266 | anthocyanidin 3-O-glucosyltransferase 2-like | | |
| tube_chr9_000836.1 | cluster271 | Fn3 like domain-containing protein | P:GO:0031222; P:GO:0045493; F:GO:0009044; F:GO:0046556 | P:arabinan catabolic process; P:xylan catabolic process; F:xylan 1,4-beta-xylosidase activity; F:alpha-L-arabinofuranosidase activity |
| tube_chr9_000835.1 | cluster271 | Fn3 like domain-containing protein | P:GO:0031222; P:GO:0045493; F:GO:0009044; F:GO:0046556 | P:arabinan catabolic process; P:xylan catabolic process; F:xylan 1,4-beta-xylosidase activity; F:alpha-L-arabinofuranosidase activity |
| tube_chr9_000834.1 | cluster271 | Fn3 like domain-containing protein | P:GO:0031222; P:GO:0045493; F:GO:0009044; F:GO:0046556 | P:arabinan catabolic process; P:xylan catabolic process; F:xylan 1,4-beta-xylosidase activity; F:alpha-L-arabinofuranosidase activity |
| tube_chr9_000509.1 | cluster271 | probable beta-D-xylosidase 2 | P:GO:0031222; P:GO:0045493; F:GO:0009044; F:GO:0046556 | P:arabinan catabolic process; P:xylan catabolic process; F:xylan 1,4-beta-xylosidase activity; F:alpha-L-arabinofuranosidase activity |
| tube_chr16_000455.1 | cluster271 | Fn3 like domain-containing protein | P:GO:0031222; P:GO:0045493; F:GO:0009044; F:GO:0046556 | P:arabinan catabolic process; P:xylan catabolic process; F:xylan 1,4-beta-xylosidase activity; F:alpha-L-arabinofuranosidase activity |
| tube_chr16_000454.1 | cluster271 | Fn3 like domain-containing protein | P:GO:0031222; P:GO:0045493; F:GO:0009044; F:GO:0046556 | P:arabinan catabolic process; P:xylan catabolic process; F:xylan 1,4-beta-xylosidase activity; F:alpha-L-arabinofuranosidase activity |
| tube_chr14_000379.1 | cluster276 | Receptor-like protein 14 | C:GO:0005886; C:GO:0016020 | C:plasma membrane; C:membrane |
| tube_chr14_000361.1 | cluster276 | Receptor-like protein 15 | C:GO:0016020 | C:membrane |
| tube_chr14_000364.1 | cluster276 | Receptor-like protein 15 | C:GO:0016020 | C:membrane |
| tube_chr14_000367.1 | cluster276 | Receptor-like protein 15 | C:GO:0016020 | C:membrane |
| tube_chr14_000371.1 | cluster276 | Receptor-like protein 15 | C:GO:0016020 | C:membrane |
| tube_chr14_000375.1 | cluster276 | Receptor-like protein 15 | C:GO:0016020 | C:membrane |
| tube_chr14_000376.1 | cluster276 | Receptor-like protein 1 | C:GO:0005886; C:GO:0016020 | C:plasma membrane; C:membrane |
| tube_chr14_000378.1 | cluster276 | Receptor-like protein 15 | C:GO:0016020 | C:membrane |

| Gene ID | Cluster | Function | GO IDs | GO Names |
|---|---|---|---|---|
| tube_chr14_000507.1 | cluster276 | Receptor-like protein 15 | C:GO:0016020 | C:membrane |
| tube_chr14_000377.1 | cluster276 | Receptor-like protein 15 | C:GO:0016020 | C:membrane |
| tube_chr5_000101.1 | cluster280 | hypothetical protein Ddye_018184 | C:GO:0016020 | C:membrane |
| tube_chr5_000102.1 | cluster280 | hypothetical protein Dsin_026442 | C:GO:0016020 | C:membrane |
| tube_chr5_000103.1 | cluster280 | hypothetical protein Dsin_026442 | C:GO:0016020 | C:membrane |
| tube_chr5_001324.1 | cluster280 | hypothetical protein Patl1_19986 | C:GO:0016020 | C:membrane |
| tube_chr5_001325.1 | cluster280 | hypothetical protein Dsin_026442 | C:GO:0016020 | C:membrane |
| tube_chr5_001326.1 | cluster280 | hypothetical protein Patl1_19986 | C:GO:0016020 | C:membrane |
| tube_chr3_001046.1 | cluster333 | probable mannitol dehydrogenase | P:GO:0009809; F:GO:0008270; F:GO:0045551 | P:lignin biosynthetic process; F:zinc ion binding; F:cinnamyl-alcohol dehydrogenase activity |
| tube_chr3_001047.1 | cluster333 | probable mannitol dehydrogenase | P:GO:0009809; F:GO:0008270; F:GO:0045551 | P:lignin biosynthetic process; F:zinc ion binding; F:cinnamyl-alcohol dehydrogenase activity |
| tube_chr3_001050.1 | cluster333 | probable mannitol dehydrogenase | P:GO:0009809; F:GO:0008270; F:GO:0045551 | P:lignin biosynthetic process; F:zinc ion binding; F:cinnamyl-alcohol dehydrogenase activity |
| tube_chr3_001048.1 | cluster333 | probable mannitol dehydrogenase | P:GO:0009809; F:GO:0008270; F:GO:0045551 | P:lignin biosynthetic process; F:zinc ion binding; F:cinnamyl-alcohol dehydrogenase activity |
| tube_chr1_000195.1 | cluster362 | (+)-neomenthol dehydrogenase-like | F:GO:0016616; C:GO:0016020 | F:oxidoreductase activity, acting on the CH-OH group of donors, NAD or NADP as acceptor; C:membrane |
| tube_chr1_000192.1 | cluster362 | (+)-neomenthol dehydrogenase-like | F:GO:0016616; C:GO:0016020 | F:oxidoreductase activity, acting on the CH-OH group of donors, NAD or NADP as acceptor; C:membrane |
| tube_chr1_002927.1 | cluster362 | (+)-neomenthol dehydrogenase-like | F:GO:0016616; C:GO:0016020 | F:oxidoreductase activity, acting on the CH-OH group of donors, NAD or NADP as acceptor; C:membrane |
| tube_chr1_000193.1 | cluster362 | (+)-neomenthol dehydrogenase-like | F:GO:0016616; C:GO:0016020 | F:oxidoreductase activity, acting on the CH-OH group of donors, NAD or NADP as acceptor; C:membrane |
| tube_chr1_000194.1 | cluster362 | (+)-neomenthol dehydrogenase-like | F:GO:0016616; C:GO:0016020 | F:oxidoreductase activity, acting on the CH-OH group of donors, NAD or NADP as acceptor; C:membrane |
| tube_chr1_000191.1 | cluster362 | (+)-neomenthol dehydrogenase-like | F:GO:0016616; C:GO:0016020 | F:oxidoreductase activity, acting on the CH-OH group of donors, NAD or NADP as acceptor; C:membrane |
| tube_chr16_001030.1 | cluster405 | probable leucine-rich repeat receptor-like serine/threonine-protein kinase At3g14840 | P:GO:0006338; F:GO:0004676; F:GO:0004677; F:GO:0004679; F:GO:0004694; F:GO:0004711; F:GO:0005524; F:GO:0035175; F:GO:0035402; F:GO:0035403; F:GO:0035979; F:GO:0044022; F:GO:0044023; F:GO:0044024; F:GO:0044025; F:GO:0072354; F:GO:0072371; F:GO:0072518; F:GO:0140823; F:GO:0140855; F:GO:0140857; F:GO:1990244; C:GO:0016020 | P:chromatin remodeling; F:3-phosphoinositide-dependent protein kinase activity; F:DNA-dependent protein kinase activity; F:AMP-activated protein kinase activity; F:eukaryotic translation initiation factor 2alpha kinase activity; F:ribosomal protein S6 kinase activity; F:ATP binding; F:histone H3S10 kinase activity; F:histone H3T11 kinase activity; F:histone H3T6 kinase activity; F:histone H2AXS139 kinase activity; F:histone H3S28 kinase activity; F:histone H4S1 kinase activity; F:histone H2AS1 kinase activity; F:histone H2BS14 kinase activity; F:histone H3T3 kinase activity; F:histone H2AS121 kinase activity; F:Rho-dependent protein serine/threonine kinase activity; F:histone H2BS36 kinase activity; F:histone H3S57 kinase activity; F:histone H3T45 kinase activity; F:histone H2AT120 kinase activity; C:membrane |
| tube_chr16_001025.1 | cluster405 | probable leucine-rich repeat receptor-like serine/threonine-protein kinase At3g14840 isoform X1 | P:GO:0006338; F:GO:0004676; F:GO:0004677; F:GO:0004679; F:GO:0004694; F:GO:0004711; F:GO:0005524; F:GO:0033612; F:GO:0035175; F:GO:0035402; F:GO:0035403; F:GO:0035979; F:GO:0044022; F:GO:0044023; F:GO:0044024; F:GO:0044025; F:GO:0072354; F:GO:0072371; F:GO:0072518; F:GO:0140823; F:GO:0140855; F:GO:0140857; F:GO:1990244; C:GO:0016020 | P:chromatin remodeling; F:3-phosphoinositide-dependent protein kinase activity; F:DNA-dependent protein kinase activity; F:AMP-activated protein kinase activity; F:eukaryotic translation initiation factor 2alpha kinase activity; F:ribosomal protein S6 kinase activity; F:ATP binding; F:receptor serine/threonine kinase binding; F:histone H3S10 kinase activity; F:histone H3T11 kinase activity; F:histone H3T6 kinase activity; F:histone H2AXS139 kinase activity; F:histone H3S28 kinase activity; F:histone H4S1 kinase activity; F:histone H2AS1 kinase activity; F:histone H2BS14 kinase activity; F:histone H3T3 kinase activity; F:histone H2AS121 kinase activity; F:Rho-dependent protein serine/threonine kinase activity; F:histone H2BS36 kinase activity; F:histone H3S57 kinase activity; F:histone H3T45 kinase activity; F:histone H2AT120 kinase activity; C:membrane |
| tube_chr16_001024.1 | cluster405 | probable leucine-rich repeat receptor-like serine/threonine-protein kinase At3g14840 isoform X1 | F:GO:0004674; F:GO:0033612; F:GO:0140995; F:GO:0140996; F:GO:0140998; C:GO:0016020 | F:protein serine/threonine kinase activity; F:receptor serine/threonine kinase binding; F:histone H2A kinase activity; F:histone H3 kinase activity; F:histone H2B kinase activity; C:membrane |
| tube_chr16_001027.1 | cluster405 | probable leucine-rich repeat receptor-like serine/threonine-protein kinase At3g14840 isoform X1 | P:GO:0006338; F:GO:0004676; F:GO:0004677; F:GO:0004679; F:GO:0004694; F:GO:0004711; F:GO:0005524; F:GO:0033612; F:GO:0035175; F:GO:0035402; F:GO:0035403; F:GO:0035979; F:GO:0044022; F:GO:0044023; F:GO:0044024; F:GO:0044025; F:GO:0072354; F:GO:0072371; F:GO:0072518; F:GO:0140823; F:GO:0140855; F:GO:0140857; F:GO:1990244; C:GO:0016020 | P:chromatin remodeling; F:3-phosphoinositide-dependent protein kinase activity; F:DNA-dependent protein kinase activity; F:AMP-activated protein kinase activity; F:eukaryotic translation initiation factor 2alpha kinase activity; F:ribosomal protein S6 kinase activity; F:ATP binding; F:receptor serine/threonine kinase binding; F:histone H3S10 kinase activity; F:histone H3T11 kinase activity; F:histone H3T6 kinase activity; F:histone H2AXS139 kinase activity; F:histone H3S28 kinase activity; F:histone H4S1 kinase activity; F:histone H2AS1 kinase activity; F:histone H2BS14 kinase activity; F:histone H3T3 kinase activity; F:histone H2AS121 kinase activity; F:Rho-dependent protein serine/threonine kinase activity; F:histone H2BS36 kinase activity; F:histone H3S57 kinase activity; F:histone H3T45 kinase activity; F:histone H2AT120 kinase activity; C:membrane |
| tube_chr5_000255.1 | cluster408 | protein PELPK1-like | | |
| tube_chr5_000259.1 | cluster408 | protein PELPK1-like | C:GO:0016020 | C:membrane |
| tube_chr5_000257.1 | cluster408 | protein PELPK1-like | C:GO:0016020 | C:membrane |
| tube_chr5_001148.1 | cluster408 | ---NA--- | | |
| tube_chr3_000059.1 | cluster418 | cytochrome P450 71AU50-like | F:GO:0004497; F:GO:0005506; F:GO:0016705; F:GO:0020037; C:GO:0016020 | F:monooxygenase activity; F:iron ion binding; F:oxidoreductase activity, acting on paired donors, with incorporation or reduction of molecular oxygen; F:heme binding; C:membrane |
| tube_chr3_000057.1 | cluster418 | cytochrome P450 71AU50-like | F:GO:0004497; F:GO:0005506; F:GO:0016705; F:GO:0020037; C:GO:0016020 | F:monooxygenase activity; F:iron ion binding; F:oxidoreductase activity, acting on paired donors, with incorporation or reduction of molecular oxygen; F:heme binding; C:membrane |

| Gene ID | Cluster | Function | GO IDs | GO Names |
|---|---|---|---|---|
| tube_chr3_000060.1 | cluster418 | cytochrome P450 71AU50-like | F:GO:0004497; F:GO:0005506; F:GO:0016705; F:GO:0020037; C:GO:0016020 | F:monooxygenase activity; F:iron ion binding; F:oxidoreductase activity, acting on paired donors, with incorporation or reduction of molecular oxygen; F:heme binding; C:membrane |
| tube_chr3_001231.1 | cluster434 | probable cysteine desulfurase | F:GO:0031071 | F:cysteine desulfurase activity |
| tube_chr3_000158.1 | cluster434 | probable cysteine desulfurase | | |
| tube_chr3_000157.1 | cluster434 | probable cysteine desulfurase | | |
| tube_chr11_000746.1 | cluster448 | glucan endo-1,3-beta-glucosidase, basic isoform-like | P:GO:0005975; F:GO:0042973 | P:carbohydrate metabolic process; F:glucan endo-1,3-beta-D-glucosidase activity |
| tube_chr11_000748.1 | cluster448 | glucan endo-1,3-beta-glucosidase, basic isoform-like | P:GO:0005975; F:GO:0042973 | P:carbohydrate metabolic process; F:glucan endo-1,3-beta-D-glucosidase activity |
| tube_chr11_000750.1 | cluster448 | glucan endo-1,3-beta-glucosidase, basic isoform-like | P:GO:0005975; F:GO:0042973 | P:carbohydrate metabolic process; F:glucan endo-1,3-beta-D-glucosidase activity |
| tube_chr11_000753.1 | cluster448 | glucan endo-1,3-beta-glucosidase, basic isoform-like | P:GO:0005975; F:GO:0042973 | P:carbohydrate metabolic process; F:glucan endo-1,3-beta-D-glucosidase activity |
| tube_chr11_000754.1 | cluster448 | glucan endo-1,3-beta-glucosidase, basic isoform-like | P:GO:0005975; F:GO:0042973 | P:carbohydrate metabolic process; F:glucan endo-1,3-beta-D-glucosidase activity |
| tube_chr10_000018.1 | cluster462 | licodione synthase-like | F:GO:0004497; F:GO:0005506; F:GO:0016705; F:GO:0020037; C:GO:0016020 | F:monooxygenase activity; F:iron ion binding; F:oxidoreductase activity, acting on paired donors, with incorporation or reduction of molecular oxygen; F:heme binding; C:membrane |
| tube_chr10_000015.1 | cluster462 | licodione synthase-like | F:GO:0004497; F:GO:0005506; F:GO:0016705; F:GO:0020037; C:GO:0016020 | F:monooxygenase activity; F:iron ion binding; F:oxidoreductase activity, acting on paired donors, with incorporation or reduction of molecular oxygen; F:heme binding; C:membrane |
| tube_chr10_000011.1 | cluster462 | licodione synthase-like | F:GO:0004497; F:GO:0005506; F:GO:0016705; F:GO:0020037; C:GO:0016020 | F:monooxygenase activity; F:iron ion binding; F:oxidoreductase activity, acting on paired donors, with incorporation or reduction of molecular oxygen; F:heme binding; C:membrane |
| tube_chr16_000752.1 | cluster503 | cytochrome P450 71D10-like | F:GO:0004497; F:GO:0005506; F:GO:0016705; F:GO:0020037 | F:monooxygenase activity; F:iron ion binding; F:oxidoreductase activity, acting on paired donors, with incorporation or reduction of molecular oxygen; F:heme binding |
| tube_chr16_000713.1 | cluster503 | cytochrome P450 71D10-like | F:GO:0004497; F:GO:0005506; F:GO:0016705; F:GO:0020037 | F:monooxygenase activity; F:iron ion binding; F:oxidoreductase activity, acting on paired donors, with incorporation or reduction of molecular oxygen; F:heme binding |
| tube_chr16_000720.1 | cluster503 | cytochrome P450 71D10-like | F:GO:0004497; F:GO:0005506; F:GO:0016705; F:GO:0020037 | F:monooxygenase activity; F:iron ion binding; F:oxidoreductase activity, acting on paired donors, with incorporation or reduction of molecular oxygen; F:heme binding |
| tube_chr15_000855.1 | cluster506 | uncharacterized protein LOC116115315 | C:GO:0016020 | C:membrane |
| tube_chr15_000854.1 | cluster506 | uncharacterized protein LOC116115315 | C:GO:0016020 | C:membrane |
| tube_chr15_000853.1 | cluster506 | hypothetical protein ACOSQ4_033239 | C:GO:0016020 | C:membrane |
| tube_chr11_000775.1 | cluster528 | lignin-forming anionic peroxidase-like | P:GO:0006979; P:GO:0042744; P:GO:0098869; F:GO:0020037; F:GO:0046872; F:GO:0140825; C:GO:0005576 | P:response to oxidative stress; P:hydrogen peroxide catabolic process; P:cellular oxidant detoxification; F:heme binding; F:metal ion binding; F:lactoperoxidase activity; C:extracellular region |
| tube_chr11_000771.1 | cluster528 | lignin-forming anionic peroxidase-like | P:GO:0006979; P:GO:0042744; P:GO:0098869; F:GO:0004601; F:GO:0020037; F:GO:0046872; C:GO:0005576 | P:response to oxidative stress; P:hydrogen peroxide catabolic process; P:cellular oxidant detoxification; F:peroxidase activity; F:heme binding; F:metal ion binding; C:extracellular region |
| tube_chr11_000773.1 | cluster528 | lignin-forming anionic peroxidase-like | P:GO:0006979; P:GO:0042744; P:GO:0098869; F:GO:0004601; F:GO:0020037; F:GO:0046872; C:GO:0005576 | P:response to oxidative stress; P:hydrogen peroxide catabolic process; P:cellular oxidant detoxification; F:peroxidase activity; F:heme binding; F:metal ion binding; C:extracellular region |
| tube_chr11_000774.1 | cluster528 | lignin-forming anionic peroxidase-like | P:GO:0006979; P:GO:0042744; P:GO:0098869; F:GO:0020037; F:GO:0046872; F:GO:0140825; C:GO:0005576 | P:response to oxidative stress; P:hydrogen peroxide catabolic process; P:cellular oxidant detoxification; F:heme binding; F:metal ion binding; F:lactoperoxidase activity; C:extracellular region |
| tube_chr5_001277.1 | cluster529 | exopolygalacturonase clone GBGE184 | P:GO:0009987; F:GO:0004553 | P:cellular process; F:hydrolase activity, hydrolyzing O-glycosyl compounds |
| tube_chr4_000374.1 | cluster529 | exopolygalacturonase clone GBGE184 | P:GO:0009987; F:GO:0004553 | P:cellular process; F:hydrolase activity, hydrolyzing O-glycosyl compounds |
| tube_chr4_000373.1 | cluster529 | exopolygalacturonase clone GBGE184 | P:GO:0009987; F:GO:0004553 | P:cellular process; F:hydrolase activity, hydrolyzing O-glycosyl compounds |
| tube_chr4_000372.1 | cluster529 | exopolygalacturonase clone GBGE184 | P:GO:0009987; F:GO:0004553 | P:cellular process; F:hydrolase activity, hydrolyzing O-glycosyl compounds |
| tube_chr4_000371.1 | cluster529 | exopolygalacturonase clone GBGE184 | P:GO:0009987; F:GO:0004553 | P:cellular process; F:hydrolase activity, hydrolyzing O-glycosyl compounds |
| tube_chr5_001276.1 | cluster529 | exopolygalacturonase clone GBGE184 | P:GO:0009987; F:GO:0004553 | P:cellular process; F:hydrolase activity, hydrolyzing O-glycosyl compounds |
| tube_chr3_001262.1 | cluster535 | monooxygenase 2-like | | |
| tube_chr3_001263.1 | cluster535 | monooxygenase 2-like | | |
| tube_chr3_001264.1 | cluster535 | monooxygenase 2-like | | |
| tube_chr3_001266.1 | cluster535 | monooxygenase 2-like | | |
| tube_chr3_001265.1 | cluster535 | monooxygenase 2-like | | |
| tube_chr3_001267.1 | cluster535 | monooxygenase 2-like | | |
| tube_chr9_001237.1 | cluster542 | ankyrin repeat-containing protein NPR4-like | C:GO:0016020 | C:membrane |

| Gene ID | Cluster | Function | GO IDs | GO Names |
|---|---|---|---|---|
| tube_chr9_001238.1 | cluster542 | ankyrin repeat-containing protein NPR4-like isoform X1 | C:GO:0016020 | C:membrane |
| tube_chr9_000106.1 | cluster542 | ankyrin repeat-containing protein NPR4-like | C:GO:0016020 | C:membrane |
| tube_chr9_000100.1 | cluster542 | ankyrin repeat-containing protein NPR4-like | C:GO:0016020 | C:membrane |
| tube_chr3_000594.1 | cluster710 | S-locus F-box protein | | |
| tube_chr14_000413.1 | cluster710 | S-locus F-box protein | | |
| tube_chr14_000597.1 | cluster710 | S-locus F-box protein | | |
| tube_chr14_000599.1 | cluster710 | S-locus F-box protein | | |
| tube_chr14_000600.1 | cluster710 | S-locus F-box protein | | |
| tube_chr3_000599.1 | cluster710 | S-locus F-box protein | | |
| tube_chr3_000802.1 | cluster710 | S-locus F-box protein | | |
| tube_chr3_000803.1 | cluster710 | S-locus F-box protein | | |
| tube_chr8_001099.1 | cluster719 | Proteoglycan 4 C-terminal part like | | |
| tube_chr8_001102.1 | cluster719 | Proteoglycan 4 C-terminal part like | | |
| tube_chr8_001100.1 | cluster719 | Proteoglycan 4 C-terminal part like | | |
| tube_chr4_000049.1 | cluster739 | 2-alkenal reductase (NADP(+)-dependent) | F:GO:0032440 | F:2-alkenal reductase [NAD(P)H] activity |
| tube_chr4_000056.1 | cluster739 | 2-alkenal reductase (NADP(+)-dependent) | F:GO:0032440 | F:2-alkenal reductase [NAD(P)H] activity |
| tube_chr4_000053.1 | cluster739 | 2-alkenal reductase (NADP(+)-dependent) | F:GO:0032440 | F:2-alkenal reductase [NAD(P)H] activity |
| tube_chr4_000051.1 | cluster739 | 2-alkenal reductase (NADP(+)-dependent) | F:GO:0032440 | F:2-alkenal reductase [NAD(P)H] activity |
| tube_chr7_000028.1 | cluster742 | 2-oxoglutarate (2OG) and Fe(II)-dependent oxygenase superfamily protein | F:GO:0046872; F:GO:0051213 | F:metal ion binding; F:dioxygenase activity |
| tube_chr7_000030.1 | cluster742 | 2-oxoglutarate (2OG) and Fe(II)-dependent oxygenase superfamily protein | F:GO:0016491; F:GO:0046872; C:GO:0016020 | F:oxidoreductase activity; F:metal ion binding; C:membrane |
| tube_chr7_000033.1 | cluster742 | 2-oxoglutarate (2OG) and Fe(II)-dependent oxygenase superfamily protein | F:GO:0046872; F:GO:0051213 | F:metal ion binding; F:dioxygenase activity |
| tube_chr7_001386.1 | cluster742 | 2-oxoglutarate (2OG) and Fe(II)-dependent oxygenase superfamily protein | F:GO:0046872; F:GO:0051213 | F:metal ion binding; F:dioxygenase activity |
| tube_chr7_001387.1 | cluster742 | 2-oxoglutarate (2OG) and Fe(II)-dependent oxygenase superfamily protein | F:GO:0016491; F:GO:0046872 | F:oxidoreductase activity; F:metal ion binding |
| tube_chr16_000223.1 | cluster752 | Non-specific lipid-transfer protein | P:GO:0006869; F:GO:0008289 | P:lipid transport; F:lipid binding |
| tube_chr16_000221.1 | cluster752 | non-specific lipid-transfer protein 11 | P:GO:0006869; F:GO:0008289 | P:lipid transport; F:lipid binding |
| tube_chr16_000222.1 | cluster752 | non-specific lipid-transfer protein A-like | P:GO:0006869; F:GO:0008289 | P:lipid transport; F:lipid binding |
| tube_chr16_000224.1 | cluster752 | non-specific lipid-transfer protein 11 | P:GO:0006869; F:GO:0008289 | P:lipid transport; F:lipid binding |
| tube_chr16_000225.1 | cluster752 | non-specific lipid-transfer protein 11 | P:GO:0006869; F:GO:0008289 | P:lipid transport; F:lipid binding |
| tube_chr16_000226.1 | cluster752 | Non-specific lipid-transfer protein | P:GO:0006869; F:GO:0008289 | P:lipid transport; F:lipid binding |
| tube_chr6_000951.1 | cluster755 | probable xyloglucan endotransglucosylase/hydrolase protein 23 | P:GO:0010411; P:GO:0042546; P:GO:0071555; F:GO:0004553; F:GO:0016762; C:GO:0048046 | P:xyloglucan metabolic process; P:cell wall biogenesis; P:cell wall organization; F:hydrolase activity, hydrolyzing O-glycosyl compounds; F:xyloglucan:xyloglucosyl transferase activity; C:apoplast |
| tube_chr6_000947.1 | cluster755 | probable xyloglucan endotransglucosylase/hydrolase protein 23 | | |
| tube_chr6_000948.1 | cluster755 | probable xyloglucan endotransglucosylase/hydrolase protein 23 | P:GO:0010411; P:GO:0042546; P:GO:0071555; F:GO:0004553; F:GO:0016762; C:GO:0048046 | P:xyloglucan metabolic process; P:cell wall biogenesis; P:cell wall organization; F:hydrolase activity, hydrolyzing O-glycosyl compounds; F:xyloglucan:xyloglucosyl transferase activity; C:apoplast |
| tube_chr6_000949.1 | cluster755 | probable xyloglucan endotransglucosylase/hydrolase protein 23 | P:GO:0010411; P:GO:0042546; P:GO:0071555; F:GO:0004553; F:GO:0016762; C:GO:0048046 | P:xyloglucan metabolic process; P:cell wall biogenesis; P:cell wall organization; F:hydrolase activity, hydrolyzing O-glycosyl compounds; F:xyloglucan:xyloglucosyl transferase activity; C:apoplast |
| tube_chr14_001034.1 | cluster815 | GDSL esterase/lipase | F:GO:0016788 | F:hydrolase activity, acting on ester bonds |
| tube_chr14_000033.1 | cluster815 | GDSL esterase/lipase 1-like | F:GO:0016298 | F:lipase activity |
| tube_chr14_001073.1 | cluster815 | GDSL esterase/lipase 1-like | F:GO:0016788 | F:hydrolase activity, acting on ester bonds |
| tube_chr14_001074.1 | cluster815 | GDSL esterase/lipase | F:GO:0016788 | F:hydrolase activity, acting on ester bonds |
| tube_chr9_001094.1 | cluster843 | (3S,6E)-nerolidol synthase 1-like | P:GO:0016102; F:GO:0000287; F:GO:0010333 | P:diterpenoid biosynthetic process; F:magnesium ion binding; F:terpene synthase activity |
| tube_chr9_001095.1 | cluster843 | (3S,6E)-nerolidol synthase 1-like | P:GO:0016102; F:GO:0000287; F:GO:0010333 | P:diterpenoid biosynthetic process; F:magnesium ion binding; F:terpene synthase activity |
| tube_chr9_000290.1 | cluster843 | (3S,6E)-nerolidol synthase 1-like | P:GO:0016102; F:GO:0000287; F:GO:0010333 | P:diterpenoid biosynthetic process; F:magnesium ion binding; F:terpene synthase activity |

| Gene ID | Cluster | Function | GO IDs | GO Names |
| --- | --- | --- | --- | --- |
| tube_chr6_000804.1 | cluster855 | NAD(P)H-quinone oxidoreductase subunit 1 | P:GO:0006412; P:GO:0009060; P:GO:1902600; F:GO:0003735; F:GO:0008137; F:GO:0046872; F:GO:0048038; F:GO:0051287; F:GO:0051539; C:GO:0005840; C:GO:0005886; C:GO:0009507; C:GO:0009579; C:GO:1990904 | P:translation; P:aerobic respiration; P:proton transmembrane transport; F:structural constituent of ribosome; F:NADH dehydrogenase (ubiquinone) activity; F:metal ion binding; F:quinone binding; F:NAD binding; F:4 iron, 4 sulfur cluster binding; C:ribosome; C:plasma membrane; C:chloroplast; C:thylakoid; C:ribonucleoprotein complex |
| tube_chr6_000802.1 | cluster855 | NADH-plastoquinone oxidoreductase subunit 7 | P:GO:0019684; F:GO:0048038; F:GO:0050136; F:GO:0051287; C:GO:0005886; C:GO:0009535 | P:photosynthesis, light reaction; F:quinone binding; F:NADH dehydrogenase (quinone) (non-electrogenic) activity; F:NAD binding; C:plasma membrane; C:chloroplast thylakoid membrane |
| tube_chr15_000542.1 | cluster855 | putative NADH dehydrogenase (quinone) | P:GO:0019684; F:GO:0016655; F:GO:0048038; F:GO:0051287; C:GO:0009535 | P:photosynthesis, light reaction; F:oxidoreductase activity, acting on NAD(P)H, quinone or similar compound as acceptor; F:quinone binding; F:NAD binding; C:chloroplast thylakoid membrane |
| tube_chr14_000974.1 | cluster870 | putative disease resistance protein RGA3 | P:GO:0006952; P:GO:0051707; F:GO:0043531 | P:defense response; P:response to other organism; F:ADP binding |
| tube_chr14_000966.1 | cluster870 | putative disease resistance protein RGA3 | | |
| tube_chr14_000970.1 | cluster870 | putative disease resistance protein RGA3 | | |
| tube_chr14_000972.1 | cluster870 | putative disease resistance protein RGA3 | P:GO:0006952; P:GO:0051707; F:GO:0043531 | P:defense response; P:response to other organism; F:ADP binding |
| tube_chr14_000973.1 | cluster870 | putative disease resistance protein RGA3 | | |
| tube_chr14_000975.1 | cluster870 | putative disease resistance protein RGA3 | P:GO:0006952; P:GO:0051707; F:GO:0043531 | P:defense response; P:response to other organism; F:ADP binding |
| tube_chr14_000459.1 | cluster870 | putative disease resistance protein RGA3 | | |
| tube_chr1_001266.1 | cluster990 | probable O-methyltransferase 3 | P:GO:0032259; F:GO:0008171; F:GO:0008757; F:GO:0046983 | P:methylation; F:O-methyltransferase activity; F:S-adenosylmethionine-dependent methyltransferase activity; F:protein dimerization activity |
| tube_chr1_001265.1 | cluster990 | probable O-methyltransferase 3 | P:GO:0032259; F:GO:0008171; F:GO:0008757; F:GO:0046983 | P:methylation; F:O-methyltransferase activity; F:S-adenosylmethionine-dependent methyltransferase activity; F:protein dimerization activity |
| tube_chr1_001267.1 | cluster990 | probable O-methyltransferase 3 | P:GO:0032259; F:GO:0008171; F:GO:0008757; F:GO:0046983 | P:methylation; F:O-methyltransferase activity; F:S-adenosylmethionine-dependent methyltransferase activity; F:protein dimerization activity |
| tube_chr1_001270.1 | cluster990 | probable O-methyltransferase 3 | P:GO:0032259; F:GO:0008171; F:GO:0008757; F:GO:0046983 | P:methylation; F:O-methyltransferase activity; F:S-adenosylmethionine-dependent methyltransferase activity; F:protein dimerization activity |
| tube_chr1_001264.1 | cluster990 | probable O-methyltransferase 3 | P:GO:0032259; F:GO:0008171; F:GO:0008757; F:GO:0046983 | P:methylation; F:O-methyltransferase activity; F:S-adenosylmethionine-dependent methyltransferase activity; F:protein dimerization activity |
| tube_chr10_000121.1 | cluster1000 | ethylene-responsive transcription factor ERF020-like | P:GO:0006355; F:GO:0003677; F:GO:0003700; C:GO:0005634 | P:regulation of DNA-templated transcription; F:DNA binding; F:DNA-binding transcription factor activity; C:nucleus |
| tube_chr10_000118.1 | cluster1000 | ethylene-responsive transcription factor ERF020-like | P:GO:0006355; F:GO:0003677; F:GO:0003700; C:GO:0005634 | P:regulation of DNA-templated transcription; F:DNA binding; F:DNA-binding transcription factor activity; C:nucleus |
| tube_chr10_000122.1 | cluster1000 | ethylene-responsive transcription factor ERF020-like | P:GO:0006355; F:GO:0003677; F:GO:0003700; C:GO:0005634 | P:regulation of DNA-templated transcription; F:DNA binding; F:DNA-binding transcription factor activity; C:nucleus |
| tube_chr10_000123.1 | cluster1000 | ethylene-responsive transcription factor ERF020-like | P:GO:0006355; F:GO:0003677; F:GO:0003700; C:GO:0005634 | P:regulation of DNA-templated transcription; F:DNA binding; F:DNA-binding transcription factor activity; C:nucleus |
| tube_chr10_000124.1 | cluster1000 | ethylene-responsive transcription factor ERF020-like | P:GO:0006355; F:GO:0003677; F:GO:0003700; C:GO:0005634 | P:regulation of DNA-templated transcription; F:DNA binding; F:DNA-binding transcription factor activity; C:nucleus |
| tube_chr5_001312.1 | cluster1008 | protein binding | F:GO:0005515 | F:protein binding |
| tube_chr5_000110.1 | cluster1008 | protein binding | F:GO:0005515 | F:protein binding |
| tube_chr8_000335.1 | cluster1010 | protein FAR1-RELATED SEQUENCE | P:GO:0006355; F:GO:0008270; C:GO:0005634 | P:regulation of DNA-templated transcription; F:zinc ion binding; C:nucleus |
| tube_chr8_000337.1 | cluster1010 | protein FAR1-RELATED SEQUENCE | P:GO:0006355; F:GO:0008270; C:GO:0005634 | P:regulation of DNA-templated transcription; F:zinc ion binding; C:nucleus |
| tube_chr8_000336.1 | cluster1010 | protein FAR1-RELATED SEQUENCE | P:GO:0006355; F:GO:0008270; C:GO:0005634 | P:regulation of DNA-templated transcription; F:zinc ion binding; C:nucleus |
| tube_chr11_000949.1 | cluster1074 | B3 domain-containing protein | F:GO:0003677; C:GO:0005634 | F:DNA binding; C:nucleus |
| tube_chr11_000282.1 | cluster1074 | B3 domain-containing protein | F:GO:0003677; C:GO:0005634 | F:DNA binding; C:nucleus |
| tube_chr11_000950.1 | cluster1074 | B3 domain-containing protein | F:GO:0003677; C:GO:0005634 | F:DNA binding; C:nucleus |
| tube_chr5_001118.1 | cluster1225 | L-type lectin-domain containing receptor kinase IX.1-like | F:GO:0004674; F:GO:0005524; C:GO:0005576; C:GO:0005886 | F:protein serine/threonine kinase activity; F:ATP binding; C:extracellular region; C:plasma membrane |
| tube_chr5_000281.1 | cluster1225 | L-type lectin-domain containing receptor kinase IX.1 | F:GO:0004674; F:GO:0005524; C:GO:0005576; C:GO:0005886 | F:protein serine/threonine kinase activity; F:ATP binding; C:extracellular region; C:plasma membrane |
| tube_chr5_000280.1 | cluster1225 | L-type lectin-domain containing receptor kinase IX.1 | F:GO:0004674; F:GO:0005524; C:GO:0005576; C:GO:0005886 | F:protein serine/threonine kinase activity; F:ATP binding; C:extracellular region; C:plasma membrane |
| tube_chr11_000541.1 | cluster1258 | Transmembrane protein | F:GO:0003676; F:GO:0008270 | F:nucleic acid binding; F:zinc ion binding |

| Gene ID | Cluster | Function | GO IDs | GO Names |
|---|---|---|---|---|
| tube_chr11_000540.1 | cluster1258 | extracellular ribonuclease LE isoform X1 | P:GO:0006401; F:GO:0003723; F:GO:0004519; F:GO:0004521; F:GO:0016829; F:GO:0033897; C:GO:0005576 | P:RNA catabolic process; F:RNA binding; F:endonuclease activity; F:RNA endonuclease activity; F:lyase activity; F:ribonuclease T2 activity; C:extracellular region |
| tube_chr11_000542.1 | cluster1258 | extracellular ribonuclease LE isoform X1 | P:GO:0006401; F:GO:0003723; F:GO:0004519; F:GO:0004521; F:GO:0016829; F:GO:0033897; C:GO:0005576 | P:RNA catabolic process; F:RNA binding; F:endonuclease activity; F:RNA endonuclease activity; F:lyase activity; F:ribonuclease T2 activity; C:extracellular region |
| tube_chr14_000601.1 | cluster1258 | ribonuclease T2 | F:GO:0003824 | F:catalytic activity |
| tube_chr14_000596.1 | cluster1258 | ribonuclease T2 | F:GO:0003824 | F:catalytic activity |
| tube_chr14_000598.1 | cluster1258 | ribonuclease T2 | F:GO:0003824 | F:catalytic activity |
| tube_chr3_000596.1 | cluster1258 | S2-ribonuclease | F:GO:0004519 | F:endonuclease activity |
| tube_chr14_000414.1 | cluster1258 | S2-ribonuclease | P:GO:0006401; F:GO:0003723; F:GO:0004519; F:GO:0004521; F:GO:0016829; F:GO:0033897; C:GO:0005576 | P:RNA catabolic process; F:RNA binding; F:endonuclease activity; F:RNA endonuclease activity; F:lyase activity; F:ribonuclease T2 activity; C:extracellular region |
| tube_chr2_000886.1 | cluster1280 | MLP-like protein 328 | P:GO:0006952 | P:defense response |
| tube_chr2_000887.1 | cluster1280 | MLP-like protein 328 | P:GO:0006952 | P:defense response |
| tube_chr2_000888.1 | cluster1280 | MLP-like protein 328 | P:GO:0006952 | P:defense response |
| tube_chr2_000889.1 | cluster1280 | MLP-like protein 328 | P:GO:0006952 | P:defense response |
| tube_chr2_000890.1 | cluster1280 | MLP-like protein 328 | P:GO:0006952 | P:defense response |
| tube_chr2_000891.1 | cluster1280 | MLP-like protein 328 | P:GO:0006952 | P:defense response |
| tube_chr4_001356.1 | cluster1280 | MLP-like protein 328 | P:GO:0006952 | P:defense response |
| tube_chr11_000749.1 | cluster1281 | protein PFC0760c-like | | |
| tube_chr11_000745.1 | cluster1281 | protein PFC0760c-like | | |
| tube_chr11_000747.1 | cluster1281 | Zinc finger, CCHC-type | | |
| tube_chr11_000752.1 | cluster1281 | Zinc finger, CCHC-type | | |
| tube_chr5_000036.1 | cluster1442 | ANK REP REGION domain-containing protein | C:GO:0005886; C:GO:0016020 | C:plasma membrane; C:membrane |
| tube_chr4_001129.1 | cluster1442 | ANK REP REGION domain-containing protein | C:GO:0005886; C:GO:0016020 | C:plasma membrane; C:membrane |
| tube_chr3_001223.1 | cluster1442 | putative Ankyrin repeat-containing protein | C:GO:0005886; C:GO:0016020 | C:plasma membrane; C:membrane |
| tube_chr10_001021.1 | cluster1526 | putative F-box protein At2g02030 | | |
| tube_chr10_001018.1 | cluster1526 | putative F-box protein At2g02030 | | |
| tube_chr10_001020.1 | cluster1526 | putative F-box protein At2g02030 | | |
| tube_chr14_000400.1 | cluster1555 | Phospholipase-like protein | | |
| tube_chr10_000943.1 | cluster1555 | Phospholipase-like protein | P:GO:0006508; F:GO:0008234; C:GO:0016020 | P:proteolysis; F:cysteine-type peptidase activity; C:membrane |
| tube_chr4_001464.1 | cluster1555 | hypothetical protein Pint_01671 | | |
| tube_chr5_001333.1 | cluster1555 | Phospholipase-like protein | | |
| tube_chr9_000193.1 | cluster1555 | ---NA--- | | |
| tube_chr12_001024.1 | cluster1555 | ---NA--- | | |
| tube_chr8_000613.1 | cluster1555 | ---NA--- | | |
| tube_chr3_001219.1 | cluster1555 | Phospholipase-like protein | P:GO:0006508; F:GO:0008234 | P:proteolysis; F:cysteine-type peptidase activity |
| tube_chr15_000495.1 | cluster1620 | cation/H(+) antiporter 15-like | P:GO:0006812; P:GO:0055085; C:GO:0110165 | P:monoatomic cation transport; P:transmembrane transport; C:cellular anatomical structure |
| tube_chr15_000496.1 | cluster1620 | cation/H(+) antiporter 15-like | P:GO:0006812; P:GO:0055085; C:GO:0110165 | P:monoatomic cation transport; P:transmembrane transport; C:cellular anatomical structure |
| tube_chr3_001039.1 | cluster1731 | putative Kinase | P:GO:0002229; P:GO:0009626; F:GO:0004674; F:GO:0005524; F:GO:0030246; C:GO:0005886 | P:defense response to oomycetes; P:plant-type hypersensitive response; F:protein serine/threonine kinase activity; F:ATP binding; F:carbohydrate binding; C:plasma membrane |
| tube_chr3_001037.1 | cluster1731 | putative Kinase | P:GO:0002229; P:GO:0009626; F:GO:0004674; F:GO:0005524; F:GO:0030246; C:GO:0005886 | P:defense response to oomycetes; P:plant-type hypersensitive response; F:protein serine/threonine kinase activity; F:ATP binding; F:carbohydrate binding; C:plasma membrane |
| tube_chr7_000618.1 | cluster2373 | glutathione S-transferase U7 | P:GO:0006749; P:GO:0009407; F:GO:0004364; C:GO:0005829 | P:glutathione metabolic process; P:toxin catabolic process; F:glutathione transferase activity; C:cytosol |
| tube_chr7_000617.1 | cluster2373 | glutathione S-transferase U7 | P:GO:0006749; P:GO:0009407; F:GO:0004364; C:GO:0005829 | P:glutathione metabolic process; P:toxin catabolic process; F:glutathione transferase activity; C:cytosol |
| tube_chr7_000623.1 | cluster2373 | glutathione S-transferase U7 | P:GO:0006749; P:GO:0009407; F:GO:0004364; C:GO:0005829 | P:glutathione metabolic process; P:toxin catabolic process; F:glutathione transferase activity; C:cytosol |

| Gene ID | Cluster | Function | GO IDs | GO Names |
|---|---|---|---|---|
| tube_chr7_001288.1 | cluster2774 | ---NA--- | | |
| tube_chr7_000144.1 | cluster2774 | ---NA--- | | |
| tube_chr7_001289.1 | cluster2774 | ---NA--- | | |
| tube_chr9_001210.1 | cluster3658 | probable zinc transporter 10 | P:GO:0071577; F:GO:0005385; F:GO:0032440; C:GO:0005886 | P:zinc ion transmembrane transport; F:zinc ion transmembrane transporter activity; F:2-alkenal reductase [NAD(P)H] activity; C:plasma membrane |
| tube_chr9_001209.1 | cluster3658 | probable zinc transporter 10 | P:GO:0071577; F:GO:0005385; C:GO:0005886 | P:zinc ion transmembrane transport; F:zinc ion transmembrane transporter activity; C:plasma membrane |
| tube_chr4_000973.1 | cluster3669 | CASP-like protein 1D1 | C:GO:0005886 | C:plasma membrane |
| tube_chr4_000972.1 | cluster3669 | CASP-like protein 1D1 | C:GO:0005886 | C:plasma membrane |
| tube_chr4_000974.1 | cluster3669 | CASP-like protein 1D1 | C:GO:0005886 | C:plasma membrane |
| tube_chr4_001435.1 | cluster3943 | hypothetical protein Pint_26117 | | |
| tube_chr4_001438.1 | cluster3943 | hypothetical protein Pint_26117 | | |
| tube_chr8_000480.1 | cluster4350 | Ribonuclease H protein | F:GO:0003676; F:GO:0004523 | F:nucleic acid binding; F:RNA-DNA hybrid ribonuclease activity |
| tube_chr8_001129.1 | cluster4350 | Ribonuclease H protein | F:GO:0140640 | F:catalytic activity, acting on a nucleic acid |
| tube_chr12_000032.1 | cluster4350 | putative ribonuclease H-like domain-containing protein | F:GO:0003676; F:GO:0004523 | F:nucleic acid binding; F:RNA-DNA hybrid ribonuclease activity |
| tube_chr13_000863.1 | cluster4968 | probable glucan endo-1,3-beta-glucosidase BG4 | P:GO:0005975; F:GO:0042973 | P:carbohydrate metabolic process; F:glucan endo-1,3-beta-D-glucosidase activity |
| tube_chr6_001178.1 | cluster4968 | probable glucan endo-1,3-beta-glucosidase BG4 | F:GO:0004553 | F:hydrolase activity, hydrolyzing O-glycosyl compounds |
| tube_chr6_001179.1 | cluster4968 | probable glucan endo-1,3-beta-glucosidase BG4 | F:GO:0004553 | F:hydrolase activity, hydrolyzing O-glycosyl compounds |
| tube_chr1_002974.1 | cluster5112 | proliferating cell nuclear antigen-like | P:GO:0006259 | P:DNA metabolic process |
| tube_chr1_002975.1 | cluster5112 | proliferating cell nuclear antigen-like | P:GO:0006260; P:GO:0006272; P:GO:0006275; P:GO:0006298; P:GO:0019985; F:GO:0003677; F:GO:0030337; C:GO:0005634; C:GO:0043626 | P:DNA replication; P:leading strand elongation; P:regulation of DNA replication; P:mismatch repair; P:translesion synthesis; F:DNA binding; F:DNA polymerase processivity factor activity; C:nucleus; C:PCNA complex |
| tube_chr1_002977.1 | cluster5112 | proliferating cell nuclear antigen-like | P:GO:0006272; P:GO:0006275; P:GO:0006298; P:GO:0019985; F:GO:0003677; F:GO:0030337; C:GO:0043626 | P:leading strand elongation; P:regulation of DNA replication; P:mismatch repair; P:translesion synthesis; F:DNA binding; F:DNA polymerase processivity factor activity; C:PCNA complex |
| tube_chr9_000004.1 | cluster6575 | 3-hydroxyisobutyryl-CoA hydrolase 1-like | P:GO:0006574; F:GO:0003860 | P:L-valine catabolic process; F:3-hydroxyisobutyryl-CoA hydrolase activity |
| tube_chr9_000003.1 | cluster6575 | 3-hydroxyisobutyryl-CoA hydrolase 1-like | P:GO:0006574; F:GO:0003860 | P:L-valine catabolic process; F:3-hydroxyisobutyryl-CoA hydrolase activity |
| tube_chr6_001645.1 | cluster6575 | 3-hydroxyisobutyryl-CoA hydrolase | P:GO:0006574; F:GO:0003860 | P:L-valine catabolic process; F:3-hydroxyisobutyryl-CoA hydrolase activity |
| tube_chr9_000005.1 | cluster6575 | 3-hydroxyisobutyryl-CoA hydrolase 1-like | P:GO:0006574; F:GO:0003860 | P:L-valine catabolic process; F:3-hydroxyisobutyryl-CoA hydrolase activity |
| tube_chr4_000035.1 | cluster7338 | Ubiquitin carboxyl-terminal hydrolase | F:GO:0005515 | F:protein binding |
| tube_chr4_000036.1 | cluster7338 | Ubiquitin carboxyl-terminal hydrolase | F:GO:0005515 | F:protein binding |
| tube_chr9_001310.1 | cluster7509 | cysteine-rich receptor-like protein kinase 10 | C:GO:0016020 | C:membrane |
| tube_chr9_001307.1 | cluster7509 | cysteine-rich receptor-like protein kinase 10 | C:GO:0016020 | C:membrane |
| tube_chr9_001312.1 | cluster7509 | cysteine-rich receptor-like protein kinase 10 | C:GO:0016020 | C:membrane |
| tube_chr9_001313.1 | cluster7509 | cysteine-rich receptor-like protein kinase 10 | C:GO:0016020 | C:membrane |
| tube_chr9_001317.1 | cluster7509 | cysteine-rich receptor-like protein kinase 10 | C:GO:0016020 | C:membrane |
| tube_chr9_001318.1 | cluster7509 | cysteine-rich receptor-like protein kinase 10 | C:GO:0016020 | C:membrane |
| tube_chr9_001319.1 | cluster7509 | cysteine-rich receptor-like protein kinase 10 | F:GO:0004674; F:GO:0005524; C:GO:0005886; C:GO:0016020 | F:protein serine/threonine kinase activity; F:ATP binding; C:plasma membrane; C:membrane |
| tube_chr3_000280.1 | cluster7510 | UDP-glycosyltransferase 91A1 | F:GO:0008194 | F:UDP-glycosyltransferase activity |
| tube_chr3_000277.1 | cluster7510 | UDP-glycosyltransferase 91A1-like | F:GO:0008194 | F:UDP-glycosyltransferase activity |
| tube_chr16_000293.1 | cluster7542 | hypothetical protein ACOSP7_030674 | F:GO:0005515 | F:protein binding |
| tube_chr16_000282.1 | cluster7542 | F-box/FBD/LRR-repeat protein At4g26340-like | F:GO:0004842; C:GO:0005783 | F:ubiquitin-protein transferase activity; C:endoplasmic reticulum |
| tube_chr8_000039.1 | cluster7862 | hypothetical protein ACOSQ2_029345 | | |
| tube_chr8_000046.1 | cluster7862 | Ubiquitin carboxyl-terminal hydrolase-related protein | P:GO:0016579; F:GO:0004843 | P:protein deubiquitination; F:cysteine-type deubiquitinase activity |
| tube_chr8_000047.1 | cluster7862 | Ubiquitin carboxyl-terminal hydrolase-related protein | P:GO:0016579; F:GO:0004843 | P:protein deubiquitination; F:cysteine-type deubiquitinase activity |
| tube_chr8_000048.1 | cluster7862 | hypothetical protein ACOSQ2_029345 | | |
| tube_chr8_001533.1 | cluster7862 | hypothetical protein Q3G72_035201 | | |
| tube_chr7_001242.1 | cluster8023 | Ubiquitin domain-containing protein | F:GO:0005515; C:GO:0005622 | F:protein binding; C:intracellular anatomical structure |

| Gene ID | Cluster | Function | GO IDs | GO Names |
|---|---|---|---|---|
| tube_chr7_001245.1 | cluster8023 | Ubiquitin domain-containing protein | F:GO:0005515; C:GO:0005622 | F:protein binding; C:intracellular anatomical structure |
| tube_chr7_001247.1 | cluster8023 | Ubiquitin domain-containing protein | F:GO:0005515; C:GO:0005622 | F:protein binding; C:intracellular anatomical structure |
| tube_chr3_000657.1 | cluster8050 | hypothetical protein Patl1_29245 | | |
| tube_chr3_000727.1 | cluster8050 | hypothetical protein Patl1_29248 | | |
| tube_chr3_000728.1 | cluster8050 | hypothetical protein Pint_28663 | | |
| tube_chr9_000416.1 | cluster8165 | abscisic acid receptor PYL11-like | P:GO:0009738; F:GO:0004864; F:GO:0010427; F:GO:0038023; C:GO:0005634; C:GO:0005737; C:GO:0005886 | P:abscisic acid-activated signaling pathway; F:protein phosphatase inhibitor activity; F:abscisic acid binding; F:signaling receptor activity; C:nucleus; C:cytoplasm; C:plasma membrane |
| tube_chr9_000417.1 | cluster8165 | abscisic acid receptor PYL12-like | P:GO:0009738; F:GO:0004864; F:GO:0010427; F:GO:0038023; C:GO:0005634; C:GO:0005737; C:GO:0005886 | P:abscisic acid-activated signaling pathway; F:protein phosphatase inhibitor activity; F:abscisic acid binding; F:signaling receptor activity; C:nucleus; C:cytoplasm; C:plasma membrane |
| tube_chr8_000071.1 | cluster9698 | aldose 1-epimerase | P:GO:0006006; P:GO:0033499; F:GO:0004034; F:GO:0030246 | P:glucose metabolic process; P:galactose catabolic process via UDP-galactose, Leloir pathway; F:aldose 1-epimerase activity; F:carbohydrate binding |
| tube_chr8_000070.1 | cluster9698 | aldose 1-epimerase-like | P:GO:0006006; P:GO:0033499; F:GO:0004034; F:GO:0030246 | P:glucose metabolic process; P:galactose catabolic process via UDP-galactose, Leloir pathway; F:aldose 1-epimerase activity; F:carbohydrate binding |
| tube_chr4_000366.1 | cluster9811 | probable glutathione S-transferase | P:GO:0006749; P:GO:0009407; P:GO:0098869; F:GO:0004364; F:GO:0045174; C:GO:0005829 | P:glutathione metabolic process; P:toxin catabolic process; P:cellular oxidant detoxification; F:glutathione transferase activity; F:glutathione dehydrogenase (ascorbate) activity; C:cytosol |
| tube_chr4_000367.1 | cluster9811 | probable glutathione S-transferase | P:GO:0006749; P:GO:0009636; F:GO:0004364; F:GO:0004462; C:GO:0005829 | P:glutathione metabolic process; P:response to toxic substance; F:glutathione transferase activity; F:lactoylglutathione lyase activity; C:cytosol |
| tube_chr14_000495.1 | cluster11659 | methanol O-anthraniloyltransferase-like | | |
| tube_chr14_000493.1 | cluster11659 | alcohol acyl transferase 1 allele RGb-like | | |
| tube_chr14_000678.1 | cluster11659 | methanol O-anthraniloyltransferase-like | | |
| tube_chr16_001074.1 | cluster11910 | F-box/LRR-repeat protein At4g14103-like isoform X1 | F:GO:0005515 | F:protein binding |
| tube_chr16_001071.1 | cluster11910 | F-box/LRR-repeat protein At4g14103-like isoform X1 | F:GO:0005515 | F:protein binding |
| tube_chr14_000568.1 | cluster12968 | photosystem II protein D1 | P:GO:0009635; P:GO:0009772; F:GO:0005506; F:GO:0010242; F:GO:0016168; F:GO:0016682; F:GO:0045156; C:GO:0009523; C:GO:0009535 | P:response to herbicide; P:photosynthetic electron transport in photosystem II; F:iron ion binding; F:oxygen evolving activity; F:chlorophyll binding; F:oxidoreductase activity, acting on diphenols and related substances as donors, oxygen as acceptor; F:obsolete electron transporter, transferring electrons within the cyclic electron transport pathway of photosynthesis activity; C:photosystem II; C:chloroplast thylakoid membrane |
| tube_chr4_001371.1 | cluster12968 | PsbA | P:GO:0009772; F:GO:0005488; F:GO:0045156; C:GO:0009523; C:GO:0009535 | P:photosynthetic electron transport in photosystem II; F:binding; F:obsolete electron transporter, transferring electrons within the cyclic electron transport pathway of photosynthesis activity; C:photosystem II; C:chloroplast thylakoid membrane |
| tube_chr14_000467.1 | cluster12968 | PsbA | P:GO:0009772; F:GO:0016491; F:GO:0045156; C:GO:0009523; C:GO:0009535 | P:photosynthetic electron transport in photosystem II; F:oxidoreductase activity; F:obsolete electron transporter, transferring electrons within the cyclic electron transport pathway of photosynthesis activity; C:photosystem II; C:chloroplast thylakoid membrane |
| tube_chr2_000922.1 | cluster12968 | PsbA | P:GO:0009772; F:GO:0010242; F:GO:0045156; C:GO:0009523; C:GO:0009535 | P:photosynthetic electron transport in photosystem II; F:oxygen evolving activity; F:obsolete electron transporter, transferring electrons within the cyclic electron transport pathway of photosynthesis activity; C:photosystem II; C:chloroplast thylakoid membrane |
| tube_chr3_001284.1 | cluster13555 | hypothetical protein Pint_27066 | | |
| tube_chr3_001281.1 | cluster13555 | hypothetical protein Pint_27066 | | |
| tube_chr3_001282.1 | cluster13555 | hypothetical protein Pint_27066 | | |
| tube_chr3_001283.1 | cluster13555 | hypothetical protein Patl1_27677 | | |
| tube_chr3_001280.1 | cluster13555 | hypothetical protein Pint_27066 | | |
| tube_chr3_001278.1 | cluster13555 | hypothetical protein Pint_27066 | | |
| tube_chr2_000725.1 | cluster14030 | putative viral movement protein | | |
| tube_chr2_000726.1 | cluster14030 | putative viral movement protein | | |
| tube_chr6_001623.1 | cluster14054 | hypothetical protein LWI29_033215 | P:GO:0006952 | P:defense response |
| tube_chr3_001372.1 | cluster14054 | thaumatin-like protein | P:GO:0006952 | P:defense response |
| tube_chr1_001464.1 | cluster14870 | Rapid ALkalinization Factor | P:GO:0007165; P:GO:0007204; P:GO:0007267; P:GO:0010082; P:GO:0019722; P:GO:0040008; F:GO:0005179; C:GO:0005576; C:GO:0048046 | P:signal transduction; P:positive regulation of cytosolic calcium ion concentration; P:cell-cell signaling; P:regulation of root meristem growth; P:calcium-mediated signaling; P:regulation of growth; F:hormone activity; C:extracellular region; C:apoplast |
| tube_chr1_001463.1 | cluster14870 | hypothetical protein Pint_22106 | P:GO:0007165; P:GO:0040008; F:GO:0005179; C:GO:0005576 | P:signal transduction; P:regulation of growth; F:hormone activity; C:extracellular region |

| Gene ID | Cluster | Function | GO IDs | GO Names |
|---|---|---|---|---|
| tube_chr11_000673.1 | cluster15630 | pectinesterase inhibitor 28-like isoform X2 | P:GO:0009827; F:GO:0004857; F:GO:0046910; C:GO:0009505 | P:plant-type cell wall modification; F:enzyme inhibitor activity; F:pectinesterase inhibitor activity; C:plant-type cell wall |
| tube_chr11_000674.1 | cluster15630 | pectinesterase inhibitor 28-like isoform X2 | P:GO:0009827; F:GO:0004857; F:GO:0046910; C:GO:0009505 | P:plant-type cell wall modification; F:enzyme inhibitor activity; F:pectinesterase inhibitor activity; C:plant-type cell wall |
| tube_chr1_002512.1 | cluster15637 | Plant self-incompatibility S1 | P:GO:0060320; C:GO:0005576 | P:rejection of self pollen; C:extracellular region |
| tube_chr1_002511.1 | cluster15637 | Plant self-incompatibility S1 | P:GO:0060320; C:GO:0005576 | P:rejection of self pollen; C:extracellular region |
| tube_chr6_000036.1 | cluster16540 | EG45-like domain containing protein | P:GO:0009627; C:GO:0016020; C:GO:0048046 | P:systemic acquired resistance; C:membrane; C:apoplast |
| tube_chr6_000034.1 | cluster16540 | EG45-like domain containing protein | P:GO:0009627; C:GO:0016020; C:GO:0048046 | P:systemic acquired resistance; C:membrane; C:apoplast |
| tube_chr6_000037.1 | cluster16540 | EG45-like domain containing protein | P:GO:0009627; C:GO:0016020; C:GO:0048046 | P:systemic acquired resistance; C:membrane; C:apoplast |
| tube_chr6_000967.1 | cluster17771 | hypothetical protein EZV62_022893 | P:GO:0008152; F:GO:0008233; F:GO:0046872; C:GO:0110165 | P:metabolic process; F:peptidase activity; F:metal ion binding; C:cellular anatomical structure |
| tube_chr11_000165.1 | cluster17771 | hypothetical protein EZV62_001103 | F:GO:0008270; C:GO:0016020 | F:zinc ion binding; C:membrane |
| tube_chr11_000098.1 | cluster18003 | RING-type domain-containing protein | P:GO:0006351; P:GO:0006357; P:GO:0051726; F:GO:0003677; C:GO:0005634; C:GO:0005654; C:GO:0017053 | P:DNA-templated transcription; P:regulation of transcription by RNA polymerase II; P:regulation of cell cycle; F:DNA binding; C:nucleus; C:nucleoplasm; C:transcription repressor complex |
| tube_chr11_000096.1 | cluster18003 | RING-type domain-containing protein | P:GO:0006351; P:GO:0006357; P:GO:0051726; F:GO:0003677; F:GO:0003723; F:GO:0008270; C:GO:0005654; C:GO:0017053 | P:DNA-templated transcription; P:regulation of transcription by RNA polymerase II; P:regulation of cell cycle; F:DNA binding; F:RNA binding; F:zinc ion binding; C:nucleoplasm; C:transcription repressor complex |
| tube_chr11_000099.1 | cluster18003 | RING-type domain-containing protein | P:GO:0006351; P:GO:0006357; P:GO:0051726; F:GO:0003677; F:GO:0003723; F:GO:0008270; C:GO:0005634; C:GO:0005654; C:GO:0017053 | P:DNA-templated transcription; P:regulation of transcription by RNA polymerase II; P:regulation of cell cycle; F:DNA binding; F:RNA binding; F:zinc ion binding; C:nucleus; C:nucleoplasm; C:transcription repressor complex |
| tube_chr11_001165.1 | cluster18003 | ---NA--- | | |
| tube_chr11_000350.1 | cluster18015 | germin-like protein subfamily 1 member 20 isoform X1 | F:GO:0030145; C:GO:0048046 | F:manganese ion binding; C:apoplast |
| tube_chr11_000932.1 | cluster18015 | hypothetical protein JRO89_XSUnG0149700 | F:GO:0004672; F:GO:0005524; F:GO:0030145; F:GO:0030553; C:GO:0048046 | F:protein kinase activity; F:ATP binding; F:manganese ion binding; F:cGMP binding; C:apoplast |
| tube_chr16_000611.1 | cluster18271 | probable amino acid permease 7 isoform X2 | P:GO:0003333; F:GO:0015171; C:GO:0016020 | P:amino acid transmembrane transport; F:amino acid transmembrane transporter activity; C:membrane |
| tube_chr16_000606.1 | cluster18271 | probable amino acid permease 7 isoform X2 | P:GO:0003333; F:GO:0015171; C:GO:0016020 | P:amino acid transmembrane transport; F:amino acid transmembrane transporter activity; C:membrane |
| tube_chr16_000458.1 | cluster18774 | receptor-like protein EIX1 | C:GO:0005886 | C:plasma membrane |
| tube_chr16_000459.1 | cluster18774 | receptor-like protein EIX2 | C:GO:0005886 | C:plasma membrane |
| tube_chr16_000786.1 | cluster18774 | receptor-like protein EIX1 | C:GO:0005886 | C:plasma membrane |
| tube_chr5_000234.1 | cluster19046 | 7-deoxyloganetin glucosyltransferase-like | F:GO:0080043; F:GO:0080044; C:GO:0005737 | F:quercetin 3-O-glucosyltransferase activity; F:quercetin 7-O-glucosyltransferase activity; C:cytoplasm |
| tube_chr5_000233.1 | cluster19046 | 7-deoxyloganetin glucosyltransferase | F:GO:0080043; F:GO:0080044; C:GO:0005737 | F:quercetin 3-O-glucosyltransferase activity; F:quercetin 7-O-glucosyltransferase activity; C:cytoplasm |
| tube_chr4_000116.1 | cluster19102 | WAT1-related protein At1g68170-like | P:GO:0055085; F:GO:0022857; C:GO:0016020 | P:transmembrane transport; F:transmembrane transporter activity; C:membrane |
| tube_chr4_000117.1 | cluster19102 | WAT1-related protein At1g68170-like | C:GO:0005886 | C:plasma membrane |
| tube_chr4_000118.1 | cluster19102 | WAT1-related protein At1g68170-like | C:GO:0005886 | C:plasma membrane |
| tube_chr9_000737.1 | cluster19212 | non-specific lipid-transfer protein-like | P:GO:0006869; F:GO:0008289 | P:lipid transport; F:lipid binding |
| tube_chr9_000736.1 | cluster19212 | Non-specific lipid-transfer protein | P:GO:0006869; F:GO:0008289 | P:lipid transport; F:lipid binding |
| tube_chr9_000738.1 | cluster19212 | Non-specific lipid-transfer protein | P:GO:0006869; F:GO:0008289 | P:lipid transport; F:lipid binding |
| tube_chr13_000153.1 | cluster19218 | hypothetical protein Pint_19384 | P:GO:0006355; F:GO:0003677; F:GO:0003700; C:GO:0005634 | P:regulation of DNA-templated transcription; F:DNA binding; F:DNA-binding transcription factor activity; C:nucleus |
| tube_chr5_001391.1 | cluster19218 | hypothetical protein Pint_19384 | P:GO:0006355; F:GO:0003677; F:GO:0003700; C:GO:0005634 | P:regulation of DNA-templated transcription; F:DNA binding; F:DNA-binding transcription factor activity; C:nucleus |
| tube_chr5_001392.1 | cluster19218 | hypothetical protein Pint_19384 | P:GO:0006355; F:GO:0003677; F:GO:0003700; C:GO:0005634 | P:regulation of DNA-templated transcription; F:DNA binding; F:DNA-binding transcription factor activity; C:nucleus |

### Figure S1

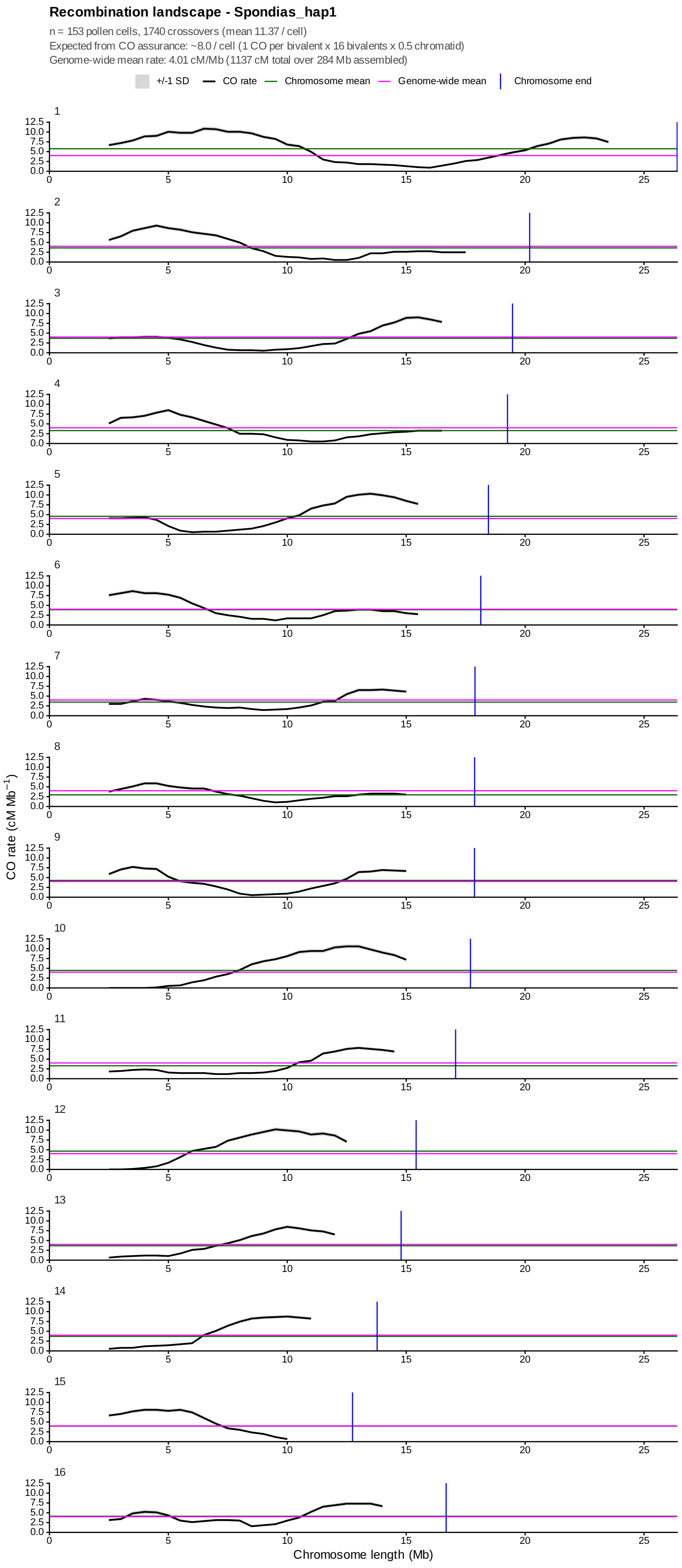

Recombination landscape - Spondias_hap1
n = 153 pollen cells, 1740 crossovers (mean 11.37 / cell)
Expected from CO assurance: ~8.0 / cell (1 CO per bivalent x 16 bivalents x 0.5 chromatid)
Genome-wide mean rate: 4.01 cM/Mb (1137 cM total over 284 Mb assembled)
+/-1 SD
CO rate
Chromosome mean
Genome-wide mean
Chromosome end
CO rate (cM Mb$^{-1}$)
Chromosome length (Mb)

**Figure S2**

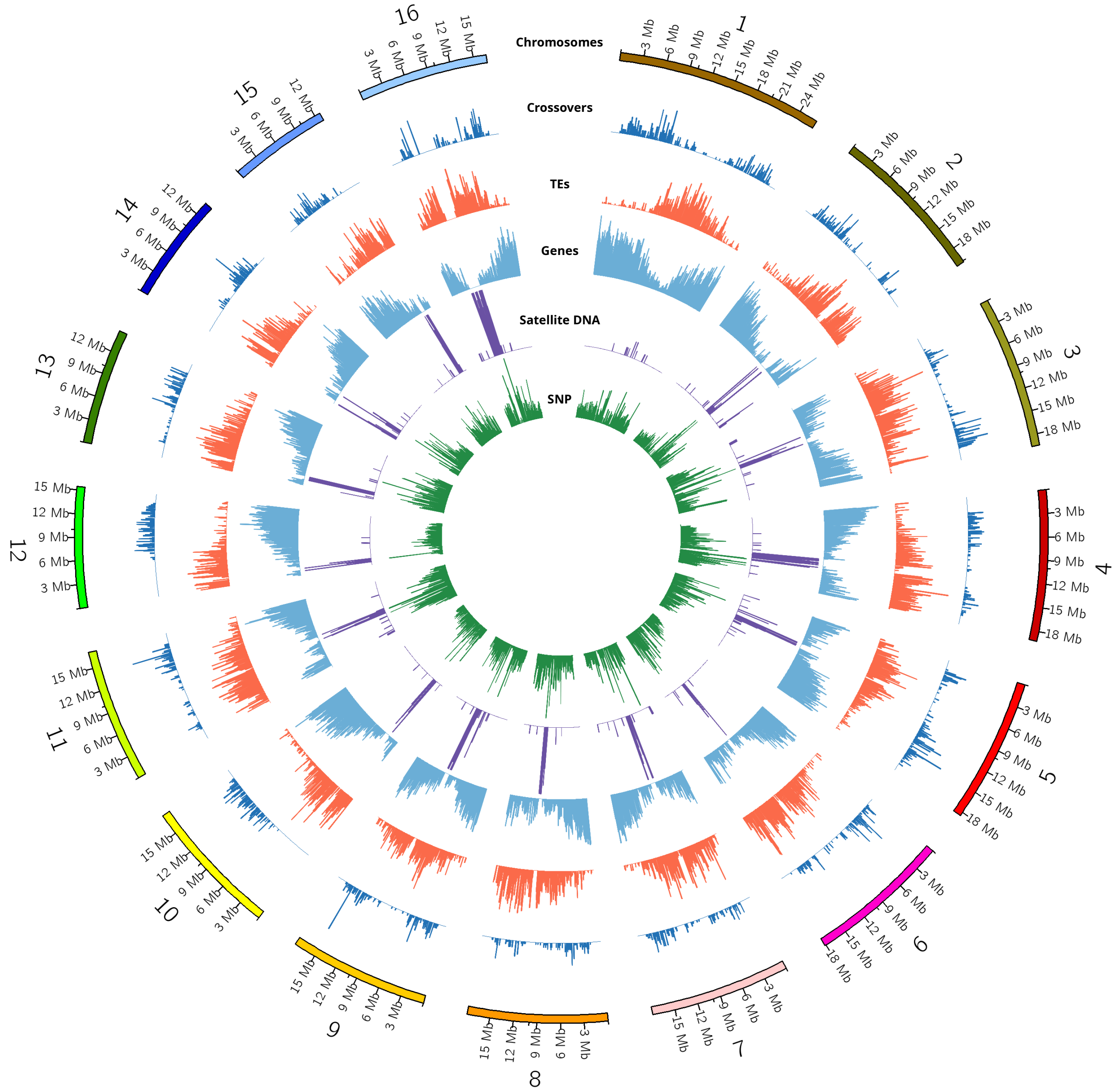
Chromosomes
Crossovers
TEs
Genes
Satellite DNA
SNP

**Figure S3**

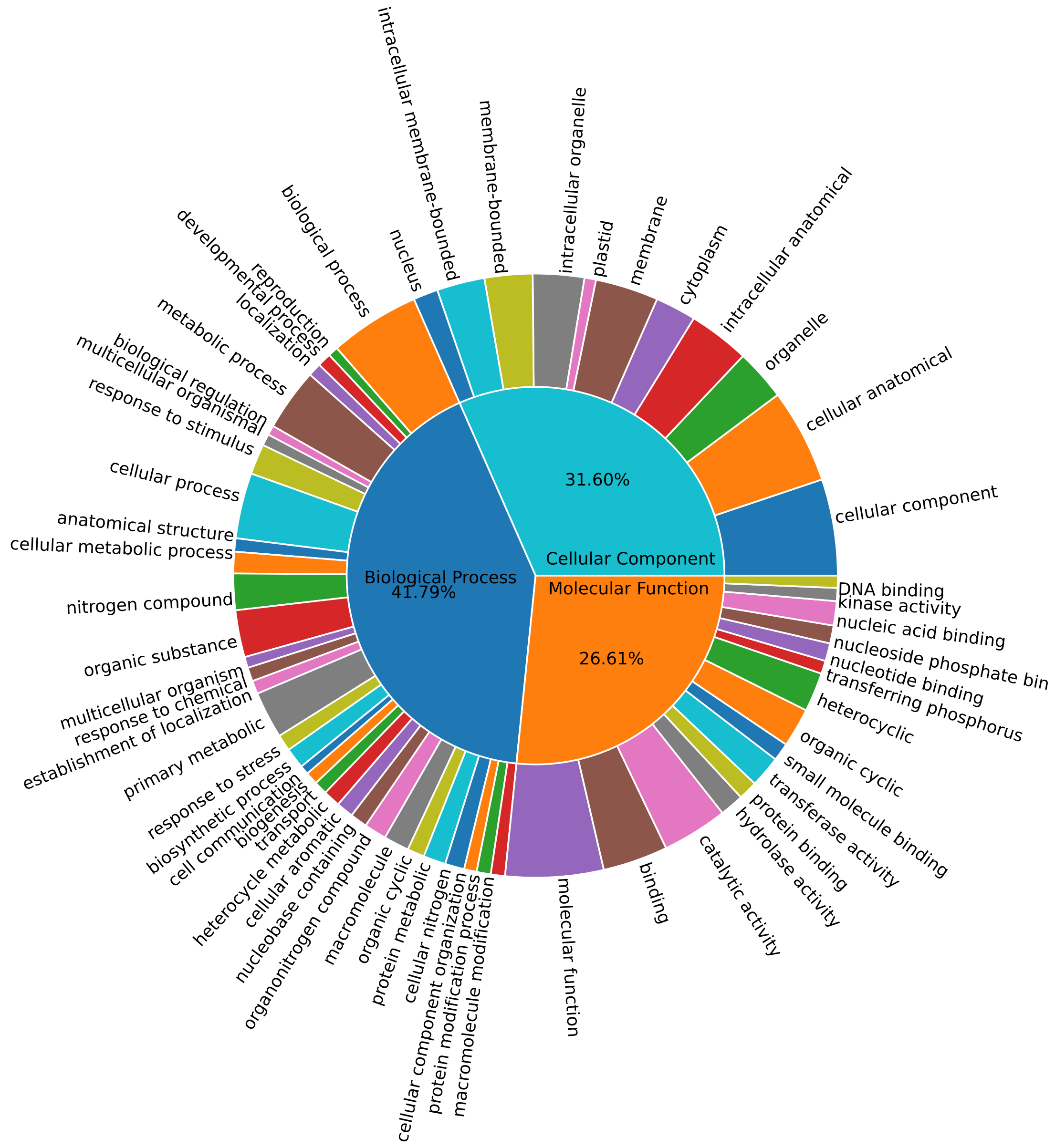

Biological Process
41.79%
Cellular Component
31.60%
Molecular Function
26.61%
cellular component
cellular anatomical
organelle
intracellular anatomical
cytoplasm
membrane
plastid
intracellular organelle
membrane-bounded
intracellular membrane-bounded
nucleus
biological process
reproduction
developmental process
localization
metabolic process
biological regulation
multicellular organismal
response to stimulus
cellular process
anatomical structure
cellular metabolic process
nitrogen compound
organic substance
multicellular organism
response to chemical
establishment of localization
primary metabolic
response to stress
biosynthetic process
cell communication
biogenesis
transport
heterocycle metabolic
cellular aromatic
nucleobase containing
organonitrogen compound
macromolecule
organic cyclic
protein metabolic
cellular nitrogen
cellular component organization
protein modification process
macromolecule modification
molecular function
binding
catalytic activity
hydrolase activity
protein binding
transferase activity
small molecule binding
organic cyclic
heterocyclic
transferring phosphorus
nucleotide binding
nucleoside phosphate bin
nucleic acid binding
kinase activity
DNA binding
